# Direct neuronal reprogramming by temporal identity factors

**DOI:** 10.1101/2021.07.05.451124

**Authors:** Camille Boudreau-Pinsonneault, Awais Javed, Michel Fries, Pierre Mattar, Michel Cayouette

## Abstract

Temporal identity factors are sufficient to reprogram developmental competence of neural progenitors, but whether they could also reprogram the identity of fully differentiated cells is unknown. To address this question, we designed a conditional gene expression system combined with genetic lineage tracing that allows rapid screening of potential reprogramming factors in the mouse retina. Using this assay, we report that co-expression of the early temporal identity transcription factor Ikzf1, together with Ikzf4, another Ikaros family member, is sufficient to directly convert adult Müller glial cells into neuron-like cells in vivo, without inducing a proliferative progenitor state. scRNA-seq analysis shows that the reprogrammed cells share some transcriptional signatures with both cone photoreceptors and bipolar cells. Furthermore, we show that co-expression of Ikzf1 and Ikzf4 can reprogram mouse embryonic fibroblasts to induced neurons by remodeling chromatin and promoting a neuronal gene expression program. This work uncovers general neuronal reprogramming properties for temporal identity factors in differentiated cells, opening new opportunities for cell therapy development.

## INTRODUCTION

Cell reprogramming has revolutionized the way we conceptualize somatic cell states. Previously assumed to be irreversible, we now know that differentiated cell type identity can be altered. Building on decades of work in *Xenopus* (Gurdon, 1962; Gurdon and Melton, 2008), Takahashi and Yamanaka (Takahashi and Yamanaka, 2006) demonstrated that the combined expression of four transcription factors mediating pluripotency during development (Oct3/4, Sox2, Klf4, c-Myc) reprograms mouse somatic cells to pluripotent cells, which can then differentiate into any cell type. Lineage-specifying transcription factors were also found to convert somatic cells from one identity to another, but without promoting an intermediate pluripotent state. For instance, the bHLH protein MyoD, a transcription factor regulating muscle cell differentiation, can reprogram mouse embryonic fibroblasts (MEFs) directly into myoblasts (Davis et al., 1987). Similarly, a small-scale screen of neural lineage transcription factors has shown that combined expression of Brn2, Ascl1, and Myt1l directly reprograms MEFs to induced neurons (iNs), wherein Ascl1 is sufficient for the conversion, and Brn2 and Myt1l potentiate neuronal differentiation (Chanda et al., 2014; Mall et al., 2017; Vierbuchen et al., 2010; Wapinski et al., 2013).

Somatic cell reprogramming offers new opportunities for tissue repair. Endogenous reprogramming is particularly appealing as it could harness the regenerative potential of resident cells, circumventing the need for difficult transplantation procedures and the low integration rate of grafted cells in host tissues (Barker et al., 2018; Li and Chen, 2016). In the central nervous system (CNS), glial cells have been targeted for endogenous reprogramming because they are present in large numbers, and are generally spared in neurodegenerative diseases. Upon injury, some mammalian glia proliferate, but undergo a wound-healing response called reactive gliosis, which is sometimes maladaptive (Bringmann et al., 2009). In contrast, the glia of non-mammalian vertebrates de-differentiate into bona fide neural progenitors capable of regenerating lost neurons (Alunni and Bally-Cuif, 2016). An extensively studied system in lower vertebrates for regenerative response in the CNS is the teleost fish retina, in which Müller glia (MG) acquire stem cell properties and regenerate retinal neurons after injury (Goldman, 2014; Lenkowski and Raymond, 2014). In contrast, the mammalian retina lacks any spontaneous regenerative response. Nonetheless, recent work in the mouse retina has identified a number of factors that can induce reprogramming of mature MG into neuron-like cells (Hoang et al., 2020; Jorstad et al., 2017; Yao et al., 2018; Zhou et al., 2020). Despite the potential of cell reprogramming for regenerative therapies in the CNS, the number of factors with the ability to reprogram fully differentiated somatic cells into therapeutically relevant neurons remains scarce, which is slowing translation efforts.

As neuronal reprogramming factors identified to date are generally involved in progenitor cell fate decisions, developmental regulators represent good candidates to identify factors with reprogramming abilities. More specifically, the functional properties of temporal identity factors, which control neural progenitor competence, make them particularly attractive. In most regions of the developing CNS, such as the neocortex, hindbrain, spinal cord, and retina, neural progenitors are temporally patterned, changing their competence to generate different cell types at specific stages (Holguera and Desplan, 2018; Oberst et al., 2019; Rossi et al., 2017). Such temporal patterning is an evolutionary conserved strategy to regulate CNS progenitor output and has been extensively studied in *Drosophila* neuroblasts. In multiple lineages of the fly ventral nerve cord, for example, a cascade of so called ‘temporal identity factors’ consisting of *hunchback (hb)*, *kruppel (kr)*, *pdm*, *castor (cas)*, and *grainyhead (grh)*, specify cell fate output and regulate temporal progression of the neuroblasts by cross-regulatory mechanisms (Brody and Odenwald, 2000; Grosskortenhaus et al., 2005; Grosskortenhaus et al., 2006; Isshiki et al., 2001; Kambadur et al., 1998; Pearson and Doe, 2003). Mis-expression of a given temporal factor in neuroblasts outside their normal window of expression is sufficient to confer competence to generate cell types associated with the temporal window of the mis-expressed factor (Cleary and Doe, 2006; Novotny et al., 2002; Pearson and Doe, 2003). In vertebrates, temporal patterning is more intricate, with complex intrinsic and extrinsic cues contributing to temporal progression of neural progenitors (Oberst et al., 2019). Interestingly, vertebrate homologues of *Drosophila* temporal identity factors have been reported to participate in temporal patterning of retinal and cortical progenitors (Alsio et al., 2013; Elliott et al., 2008; Javed et al., 2020; Mattar et al., 2015). For instance, *Ikzf1 (a.k.a. Ikaros)*, the homolog of *hb*, confers early progenitor identity, and is sufficient to induce ectopic production of early-born cell types when mis-expressed in late-stage retinal or cortical progenitor cells (Alsio et al., 2013; Elliott et al., 2008); a process that can be conceptualized as ‘temporal reprogramming’.

Although vertebrate temporal identity factors can reprogram the developmental competence of neural progenitors, it remains unknown whether they can also reprogram fully differentiated cells, and whether temporal reprogramming shares common mechanisms with other forms of reprogramming. Here we addressed these questions using the mouse retina and cultured fibroblasts as tractable systems to study cell reprogramming. Our results identify novel properties for temporal identity factors and bridge the concepts of temporal competence and cell state reprogramming.

## RESULTS

### Establishment of a screening assay for Müller glia reprogramming ex vivo

To identify new factors that can reprogram retinal glia to neurons, we designed an assay to conditionally express any gene of interest specifically in MG, while genetically tracing the lineage of the potential progenies. Although AAV vectors with GFAP mini-promoters have been used for that purpose, we decided against this approach, as leaky neuronal expression is sometimes observed with such vectors (Lee et al., 2008; Wang et al., 2020), and their specificity of expression varies depending on the transgene they carry (Su et al., 2004) (discussed in (Blackshaw and Sanes, 2021; Martin and Poche, 2019)). Instead, we relied on electroporation of conditional expression constructs (pCALM) in which a loxP-mCherry-STOP-loxP cassette is excised in a Cre-dependent manner, allowing expression of a downstream gene of interest (Fig. S1). We tested the specificity of this construct by transfecting a version containing GFP after the loxP cassette (pCALM-GFP) into HEK 293T cells, alone or together with a Cre-expressing construct. We found that only mCherry is expressed when pCALM-GFP is transfected alone, whereas mCherry is turned off and GFP is expressed when it is co-transfected with a Cre construct (Fig. S1C), validating the conditional expression strategy.

To drive Cre expression in MG, we used the Glast-CreER mice (Nathans, 2010), which specifically express a tamoxifen-inducible Cre in MG and astrocytes in the adult retina (Lehre et al., 1997). To permanently label MG and their progeny, we crossed this line with the R26R-EYFP reporter to generate Glast-CreER;R26R-EYFP mice. Seven days after tamoxifen injection in these adult mice, we found that ∼40% of all MG (CyclinD3+) are YFP+ (Fig. S2B), indicating that not all MG are targeted after tamoxifen injection. Among the population of YFP+ cells, however, we found that over 95% express CyclinD3 and 100% express glutamine synthetase, another MG marker (Fig S2A-B). Importantly, YFP+ cells were not observed in Glast-CreER;R26R-EYFP animals that did not receive tamoxifen (Fig S2C). These results show that the Glast-CreER mouse line allows specific tamoxifen-dependent activation of Cre in adult MG.

To express genes of interest specifically in MG while genetically tracing them, we electroporated pCALM constructs, individually or in combination to allow co-expression of multiple genes, in postnatal day 0-1 (P0-P1) Glast-CreER;R26R-EYFP retinas, cultured them as explants for 12 days to allow neurogenesis to complete, and then added hydroxytamoxifen in the culture medium (Fig. 1A, Fig. S1A,B). When electroporating retinas at these neonatal stages, the great majority of transfected cells are progenitors (Matsuda and Cepko, 2004), and approximately 5-10% of these progenitors generate MG (Javed et al., 2020; Matsuda and Cepko, 2004; Mattar et al., 2021), which would therefore inherit the transfected constructs (Fig. S1B). Two weeks after hydroxytamoxifen addition in the culture medium, and thus expression of the gene(s) of interest, retinas are fixed and analyzed to detect any MG cell state changes by examining morphology, soma position within the retinal layers, and expression of cell type specific markers (Fig. 1A, Fig. S1A). This assay provides a flexible and rapid way to screen for potential reprogramming effects of various genes in MG in situ.

**Figure 1:**
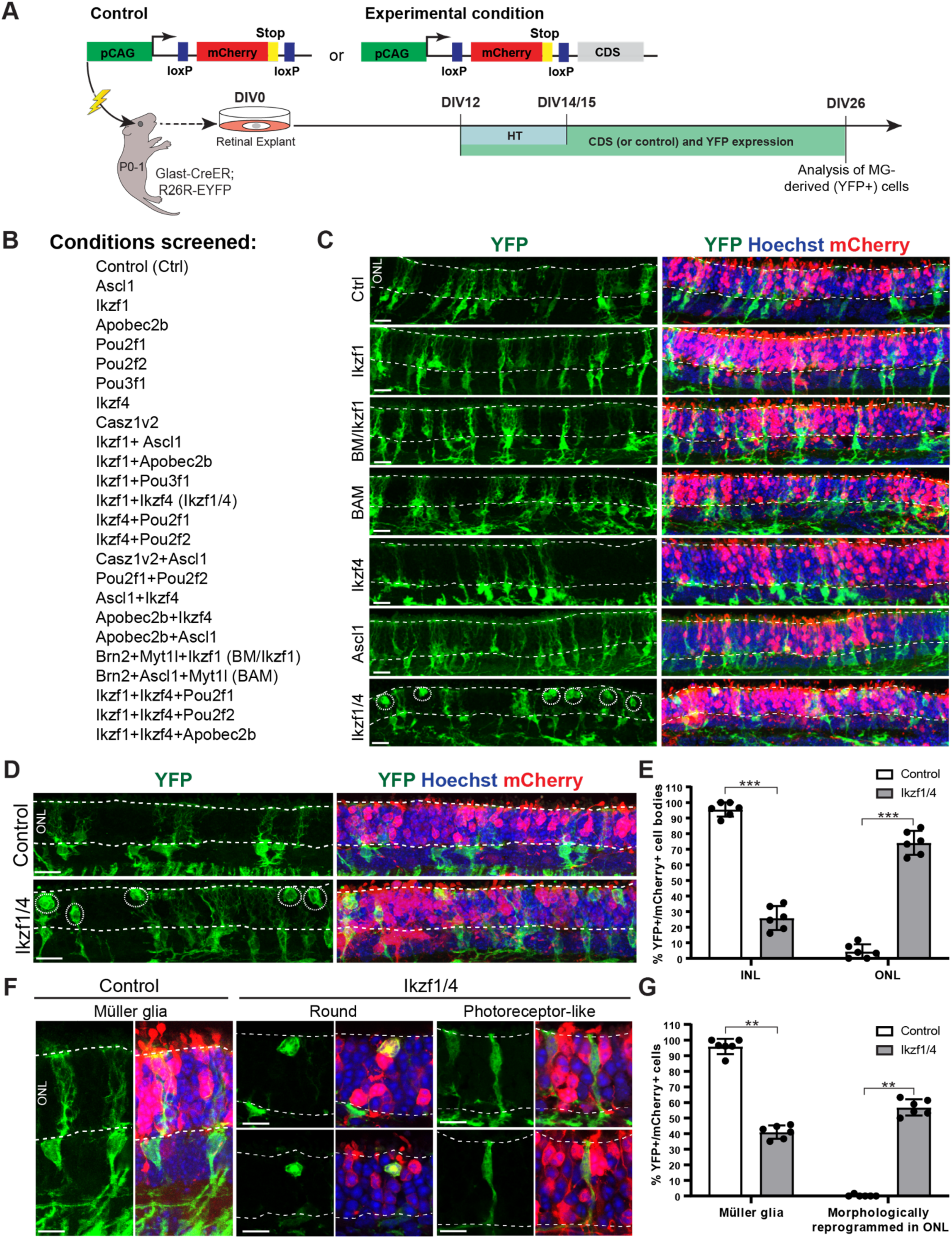
Ikzf1/4 expression induces morphological reprogramming of MG ex vivo. **(A)** Summary diagram of ex vivo screen protocol. Lightning bolt represents electroporation. HT: hydroxytamoxifen. CDS: coding sequence. **(B)** List of conditions screened for MG reprogramming. **(C)** Representative images of retinal sections electroporated with various conditions included in the screen immunostained for YFP. Dotted lines show the ONL, where photoreceptors reside. Circles in the Ikzf1/4 condition point to cells with altered morphology located in the ONL. Scale bars: 17µm. **(D)** Representative image of retinal sections from electroporated regions with control (empty pCALM vector) and Ikzf1/4 constructs immunostained for YFP. Circles point to MG-derived cells in the ONL. Dotted lines show the ONL. Scale bars: 38µm. ONL: Outer nuclear layer/photoreceptor layer **(E)** Localization of cell bodies of YFP+/mCherry+ cells for control (n=6) and Ikzf1/4 (n=6) conditions. ***p<0.001, unpaired t-test. **(F)** Representative images, immunostained for YFP, of YFP+ cells in control electroporation showing MG morphologies, and in Ikzf1/4 electroporation with rounded and photoreceptor-like morphologies. Dotted lines show the ONL. Scale bars: 10µm. **(G)** Quantifications of morphologies of control and Ikzf1/4 YFP+/mCherry+ cells. ** p=0.0022, Mann-Whitney test, control (n=6) and Ikzf1/4 (n=6). All images are z projections, except for (F) Ikzf1/4 images are single planes. Graphs represented as means +/- standard deviation.

### Combined expression of Ikzf1 and Ikzf4 reprograms Müller glia to cone-like cells ex vivo

We tested expression of 22 different gene combinations in MG (Fig. 1B), primarily focusing on various temporal identity factors previously shown to control retinal progenitor cell competence, such as Ikzf1, Pou2f1, Pou2f2, and Casz1 (Elliott et al., 2008; Javed et al., 2020; Mattar et al., 2015). Taking advantage of the flexibility of the assay, we also tested several other factors, including another Ikaros family member (Ikzf4) expressed in the retina (Clark et al., 2019; Elliott et al., 2008), factors found to promote MG reprogramming in fish (Apobec2b) (Powell et al., 2012) and mice (Ascl1) (Ueki et al., 2015), or found to potentiate MEF reprogramming (Brn2 and Myt1l) (Vierbuchen et al., 2010). In 20 out of 22 combinations tested, YFP+ cells had typical MG morphologies that were indistinguishable from those observed after expression of the empty pCALM construct, with cell bodies located in the inner nuclear layer and complex processes extending apically and basally (Fig. 1C). Importantly, these cells also expressed standard MG markers (Fig. S3A), indicating that they did not change identity. In contrast, upon Ascl1 expression, we found YFP+ cells in the inner nuclear layer that also expressed the bipolar cell marker Otx2, suggesting that some MG reprogram to bipolar-like cells, as previously reported (Pollak et al., 2013) (Fig. S3B). However, the only condition that elicited widespread changes in this assay was the combined expression of Ikzf1 and Ikzf4 (Ikzf1/4), in which many YFP+ cells with morphologies distinct from MG were observed in the outer nuclear layer, where only photoreceptor cells reside (Fig. 1C-E). More specifically, YFP+ cells were located at the apical side of the photoreceptor layer, where cone photoreceptors are found (Fig. S3C). Most relocated YFP+ cells had a round morphology without apparent cell processes, but some resembled immature photoreceptors with an oval cell body, a simple process extending towards the plexiform layer, and a small protrusion towards the apical surface (Fig. 1 F-G). Interestingly, Ikzf1 or Ikzf4 expression alone did not alter MG morphology or cell soma position (Fig. 1C), indicating that combined expression of Ikzf1 and Ikzf4 is necessary to induce morphological reprogramming. To possibly enhance Ikzf1/4 reprogramming, we also tested the co-expression of Ikzf1/4 together with other factors (Pou2f1, Pou2f2, or Apobec2b), by transfecting three constructs (Fig. 1B). We found, however, that triple transfections reduced the number of reprogrammed cells, potentially due to the dilution of each transfected construct and a reduction of the number of cells inheriting both Ikzf1 and Ikzf4. Of note, we found that the majority of morphologically-reprogrammed YFP+ cells are still mCherry+, most likely due to Cre-mediated recombination of a fraction of all plasmid constructs transfected in each cell. We hence took advantage of mCherry to focus our analysis on transfected cells for both Ikzf1/4 and control conditions (YFP+: MG and their progeny; mCherry+: transfected cells). As expected, mCherry+ MG (YFP+) co-expressed Ikzf1 and Ikzf4 after tamoxifen injection in Ikzf1/4 transfected retinas (Fig. S3D). We also validated that Ikzf1/4 are not expressed in transfected MG without tamoxifen, as expected (Fig. S3E).

To determine whether the morphologically reprogrammed cells actually change molecular identity, we performed immunofluorescence for various retinal cell type specific markers (see Table S1). Virtually all cells in the control condition expressed the MG markers Sox2 and Lhx2 (Fig. 2A, B), as expected. In contrast, most morphologically-reprogrammed YFP+ cells observed in the photoreceptor layer after Ikzf1/4 expression were negative for Sox2 and Lhx2 (Fig. 2A, B), suggesting that they lost glial identity. Nonetheless, a few cells that had their soma in the photoreceptor layer still expressed MG markers and may represent cells at intermediate stages of reprogramming (Fig. 2A, arrow). We also found that YFP+ reprogrammed cells after Ikzf1/4 expression do not stain for markers of rod photoreceptors or retinal interneurons (Table S1, Fig. S3F) and are negative for cleaved caspase 3, a marker of apoptosis (Fig. S3F). Remarkably, however, we observed that most YFP+ reprogrammed cells stain positive for the cone photoreceptor marker Rxrg (Fig. 2A, B), which is never observed in controls (Fig. 2B). Conversely, immunostaining for other cone markers (S-opsin, L/M-opsin, cone arrestin, PNA; Table S1) revealed that the YFP+ reprogrammed cells do not stain for multiple cone markers, suggesting incomplete differentiation. As Rxrg is also expressed in retinal ganglion cells, we asked whether the reprogrammed cells could be ganglion cells by staining for Brn3b. However, we found no Brn3b+/YFP+ cells (Fig. S3F), and since the reprogrammed cells moved to the photoreceptor layer, we conclude that these cells are not retinal ganglion cells. Together, these results indicate that expression of Ikzf1/4 in MG promotes reprogramming into cone-like photoreceptor cells in retinal explants.

**Figure 2:**
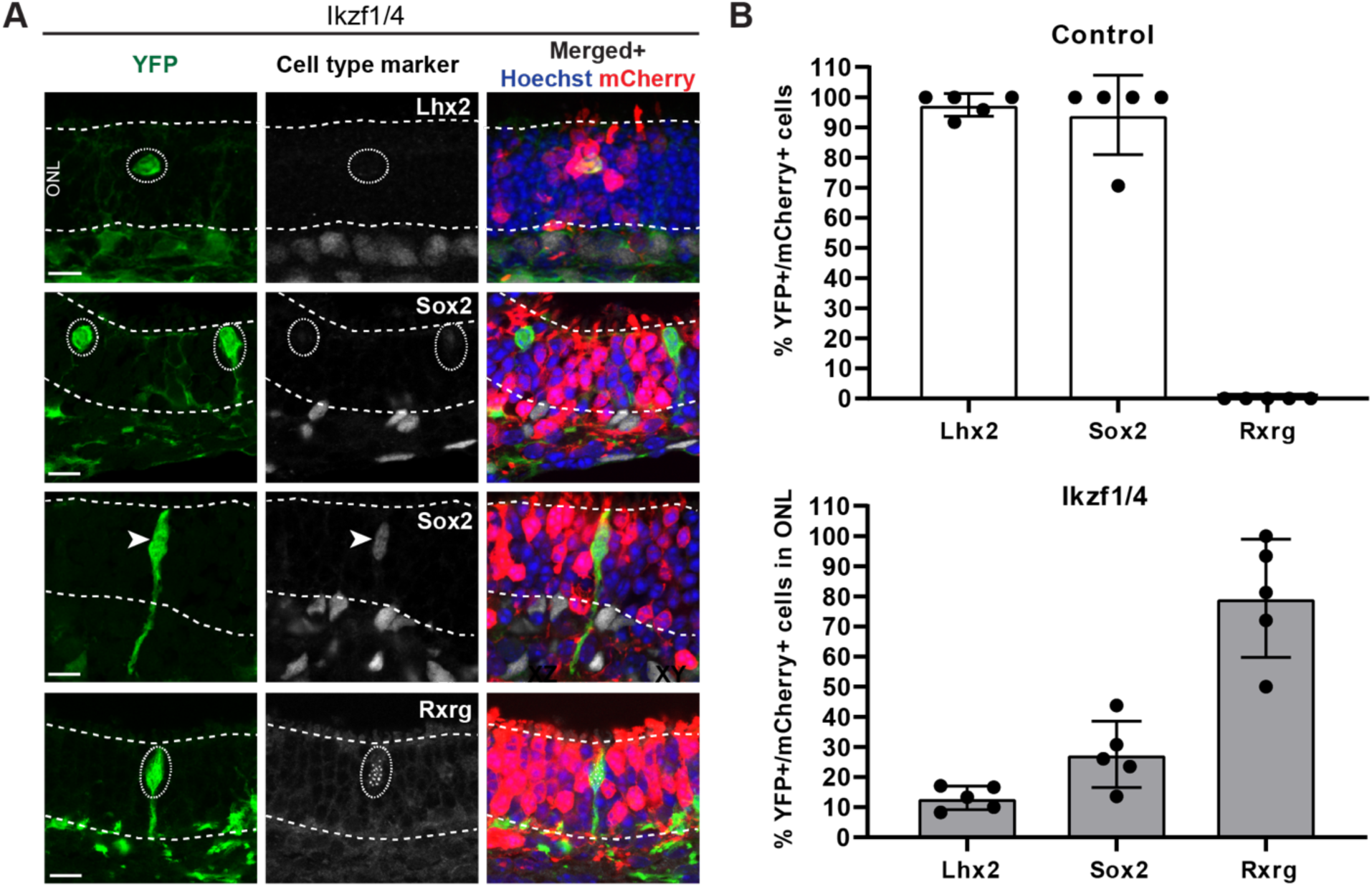
Molecular reprogramming of MG to cone-like cells following expression of Ikzf1/4 ex vivo. **(A)** Immunostaining on Ikzf1/4-electroporated retinas ex vivo with YFP and various cell type specific markers, as indicated. Most reprogrammed cells in the ONL (circled) are negative for the MG markers Lhx2 or Sox2, but are immunostained for the cone marker Rxrg. Arrow points to Sox2-positive cell in the ONL. Dotted line define the ONL thickness. Scale bars: 10µm. ONL: Outer nuclear layer/photoreceptor layer **(B)** Quantification of marker expression in control- (top) and Ikzf1/4- electroporated retinas (bottom). Images show z projections in (A) for Lhx2 and Rxrg images, all others are single planes. Graphs show means +/- standard deviation.

### Ikzf1/4 expression in Müller glia does not promote cell cycle re-entry

As in situ reprogramming in the fish retina is known to occur via de-differentiation of MG to a proliferative progenitor state (Goldman, 2014), we wondered whether Ikzf1/4-mediated reprogramming might also promote MG proliferation. To test this, we repeated the experiments described above, but added EdU to the culture medium either from DIV12 to 15 and 18 to 21 or DIV15 to 18 and 21 to 24, spanning the culture time between initiation of Ikzf1/4 expression and fixation (Fig. S4A). We found no difference between the number of YFP+/mCherry+/EdU+ cells in Ikzf1/4 compared to controls in any condition (Fig. S3B), which indicates that Ikzf1/4 do not promote MG cell cycle re-entry in this assay. Consistent with this interpretation, we found no Ki67+ cells, a proliferating progenitor marker, in the mCherry+ transfected cell population 5 days after tamoxifen administration (not shown). These results strongly suggest that Ikzf1/4-expressing mouse MG do not transition through a proliferating progenitor-like state before converting into cone-like cells, and instead are consistent with a direct transdifferentiation mechanism.

### Ikzf1/4 reprogram adult Müller glia to cone-like cells in vivo

We next investigated whether co-expression of Ikzf1/4 can also reprogram adult MG in vivo. Using a similar approach as described above in explants, we electroporated Cre-dependent empty pCALM control or conditional Ikzf1/4 expression constructs in the retinas of neonate (P0-P2) Glast-CreER;R26R-EYFP mice. When the mice reached adult ages (≥P21), we injected tamoxifen to induce YFP and Ikzf1/4 expression in MG. Three or five weeks later, the retinas were collected for analysis (Fig. 3A).

**Figure 3:**
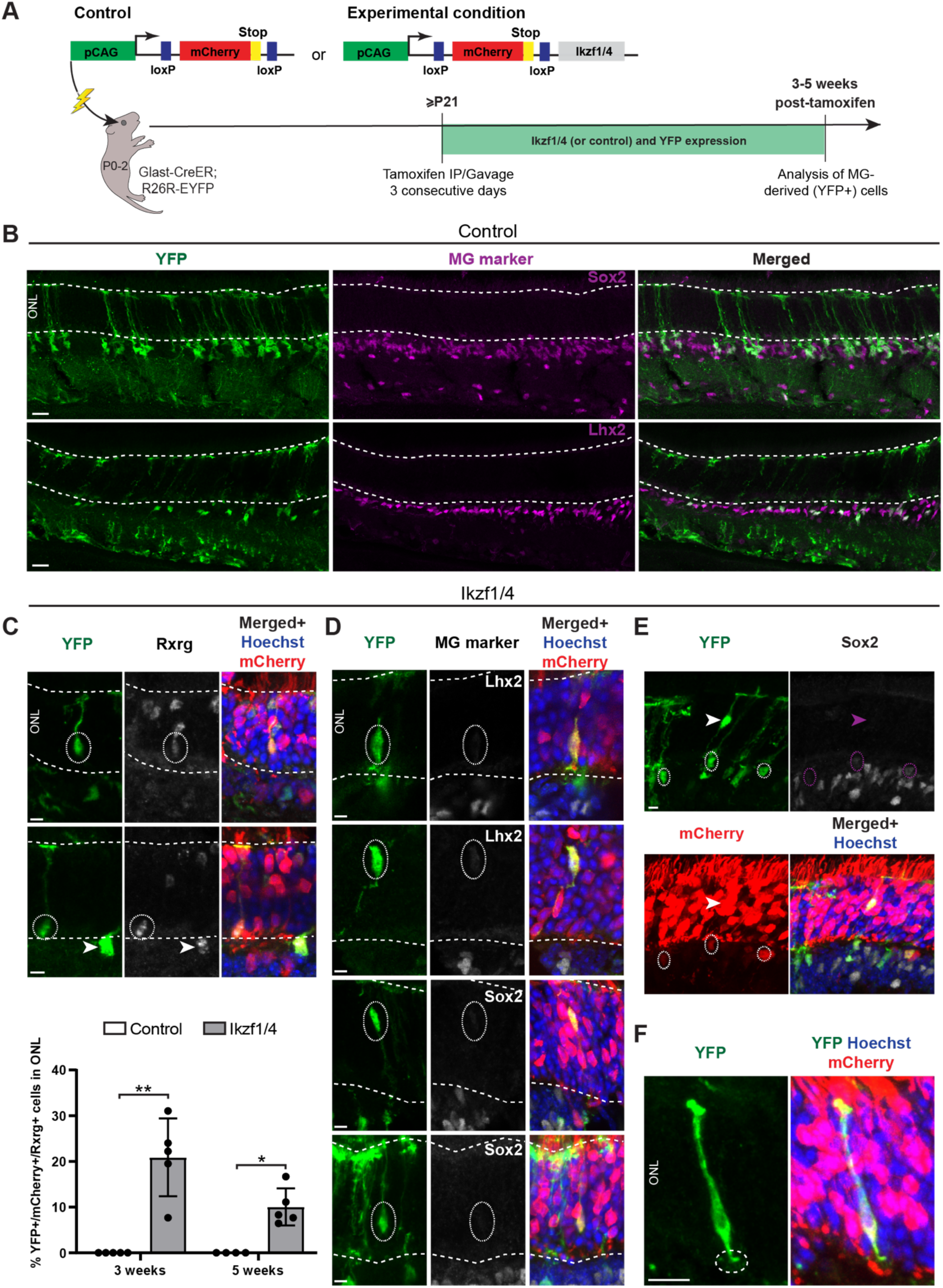
Ikzf1/4 reprogram MG to cone-like cells in vivo. **(A)** Summary diagram of the experimental protocol. IP: intraperitoneal injection. CDS: coding sequence. Lightning bolt represents electroporation. **(B)** Retinal sections from control electroporations stained for YFP and the MG markers Sox2 and Lhx2. YFP+ cells display typical MG morphology and express MG markers. Dotted lines show ONL thickness. Scale bars: 38µm. **(C)** Top: Representative images of retinal sections immunostained for YFP and the cone marker Rxrg 3 weeks after Ikzf1/4- expression in vivo. Reprogrammed cells in the ONL (circled) stain positive for Rxrg. Bottom row: Arrow shows intermediate cell in INL. Dotted lines show ONL thickness. Scale bars: 5µm. Bottom: Quantifications of reprogrammed cells in control- and Ikzf1/4-electroporated retinas at 3 and 5 weeks after tamoxifen injection. Graph represents mean +/- standard deviation. * p< 0.05; ** p<0.01; Mann-Whitney test. Control (3 weeks: n=5; 5 weeks: n=4) and Ikzf1/4 (3 weeks: n=5; 5 weeks: n=5). **(D)** Several representative images of retinal sections immunostained for YFP and the MG markers Lhx2 and Sox2 3 weeks after Ikzf1/4 expression in vivo. Ikzf1/4 reprogrammed cells in the ONL (circled) do not express MG markers. Scale bars: 5µm. **(E)** Immunostaining for YFP and Sox2 on retinal sections 3 weeks after expression of Ikzf1/4. Some YFP+/mCherry+ cells (circled) in the INL show varying levels of Sox2, whereas reprogrammed cell in the ONL (arrowhead) is negative for Sox2. Scale bar: 5µm. **(F)** High magnification image of an Ikzf1/4-reprogrammed cell, stained for YFP, showing pedicle-like structure in the outer plexiform layer (circled). Scale bars: 10µm. All images shown are z projections except for (D; top row), which is a single plane. ONL: Outer nuclear layer/photoreceptor layer; INL: Inner nuclear layer.

Three weeks post-tamoxifen, in control electroporations, we found that YFP+/mCherry+ cells have normal MG morphology with cell soma in the inner nuclear layer and expression of glia markers, as expected (Fig. 3B). After Ikzf1/4 expression, however, about a quarter of all YFP+/mCherry+ cells changed morphology, were located in the photoreceptor layer, and expressed the cone marker Rxrg (Fig. 3C). In contrast to ex vivo, we noticed that these cells do not reach the apical-most part of the photoreceptor layer, but rather span the entire thickness of this layer. The reprogrammed cells did not express the MG markers Lhx2 and Sox2 (Fig. 3D), and were still present 5 weeks post-tamoxifen, albeit in lower numbers (Fig. 3C). Remarkably, and contrary to ex vivo experiments, Ikzf1/4-reprogrammed cells very rarely showed rounded morphologies and most had complex cone-like morphologies with some pedicle-like structures in the outer plexiform layer (Fig. 3F), suggesting that the in vivo environment is more conducive to morphological differentiation. We also identified YFP+ cells in the inner nuclear layer that do not stain for the glia marker Sox2 (Fig. 3E). Some of these cells stain for Rxrg (Fig. 3C arrow), but others do not, suggesting that they might be cells in intermediate stages of reprogramming or, alternatively, reprogrammed cells of different subtypes that are not labelled by the antibodies we used. Altogether, these results show that Ikzf1/4 are sufficient to induce the conversion of adult MG into cone-like cells in vivo.

### Detection of reprogrammed cells is not the result of material exchange

Previous studies showed that transplanted photoreceptor precursor cells readily exchange cytoplasmic material, including fluorescent reporters, with host photoreceptors (Ortin-Martinez et al., 2016; Pearson et al., 2016; Santos-Ferreira et al., 2016; Singh et al., 2016), sparking the need for additional controls in lineage tracing studies with fluorescent reporters in the retina (Boudreau-Pinsonneault and Cayouette, 2018). As MG are tightly associated with cones (Reichenbach and Bringmann, 2013), we wondered whether the YFP+ cone-like cells observed after Ikzf1/4 expression might be explained by transfer of YFP from MG to endogenous cones. Our observation that reprogrammed cells do not express all cone markers and do not have mature photoreceptor morphologies argued against this interpretation, but we could not exclude the possibility that transfer of Ikzf1/4 might occur and lead to down-regulation of some cone markers. To directly address this question, we repeated the in vivo experiments, as described in Figure 3, but additionally injected the pups daily with EdU from P3 to P7 (Fig. S5A). This corresponds to the peak production period of MG from retinal progenitors, and well past the period of cone genesis. Accordingly, in control electroporations, many YFP+ MG incorporated EdU, but cone photoreceptors did not (Fig. S5B). Following expression of Ikzf1/4, we found several morphologically reprogrammed YFP+ cone-like cells in the photoreceptor layer that are also EdU+ (Fig. S5C), indicating that they are *de novo* generated cells at postnatal stages, rather than endogenous cones.

### Reprogrammed cells share transcriptional profiles with both cones and bipolars

The low throughput of immunofluorescence analyses in the above experiments rendered the precise characterization of reprogrammed cell types difficult. To circumvent this issue and obtain a more complete molecular profiling of the reprogrammed cells, we performed single cell RNA-sequencing (scRNA-seq) on sorted YFP+ cells from control- and Ikzf1/4- electroporated retinas three weeks after tamoxifen injection in adult Glast-CreER; R26R- EYFP mice (≥P21). The specificity of cell collection was tested by sorting YFP+ cells and analyzing the sorted population by flow cytometry. We found that, on average, 90% +/- 3% (n=3) of collected cells are YFP+, indicating that, although some non-fluorescent contaminating cells are present in the sorted population, we strongly enrich YFP+ cells with this method.

We sequenced 4207 cells from the control condition and 4608 cells from the Ikzf1/4 condition. As expected, the main Uniform Manifold Approximation and Projections (UMAPs) cell cluster observed in both conditions was composed of MG. Interestingly, we found a clear increase in the number of cell clusters in the Ikzf1/4 condition compared to control, and the number of cells in the MG clusters was proportionally reduced (Fig. 4A). Surprisingly, the additional clusters in the Ikzf1/4 condition were identified as bipolar cells based on top gene expression (Fig. 4B). Bipolar clusters 1, 2, 4, and 5 express markers of various bipolar cell subtypes and correspond to cone bipolar cells, whereas bipolar cluster 3 expresses rod bipolar markers (Shekhar et al., 2016) (Fig. 4C). Interestingly, we noticed that the bipolar clusters 2 and 4 additionally express *Thrb*, a cone photoreceptor gene (Fig. 4C), suggesting that these cells share transcriptional profiles with both cones and bipolars. In contrast to our immunostaining results, we did not detect Rxrg expression in these bipolar clusters, suggesting that Rxrg transcript levels may be below the detection threshold. Based on these data, we estimate that the MG to bipolar-like cell conversion rate is around 40%, which is about twice as much as the reprogramming efficiency observed by immunostaining in vivo, suggesting that a greater percentage of cells adopt a bipolar/cone transcriptomic identity than what we observed by immunostaining. Of note, we did not find progenitor-like clusters in the Ikzf1/4 condition, suggesting direct transdifferentiation of MG.

**Figure 4:**
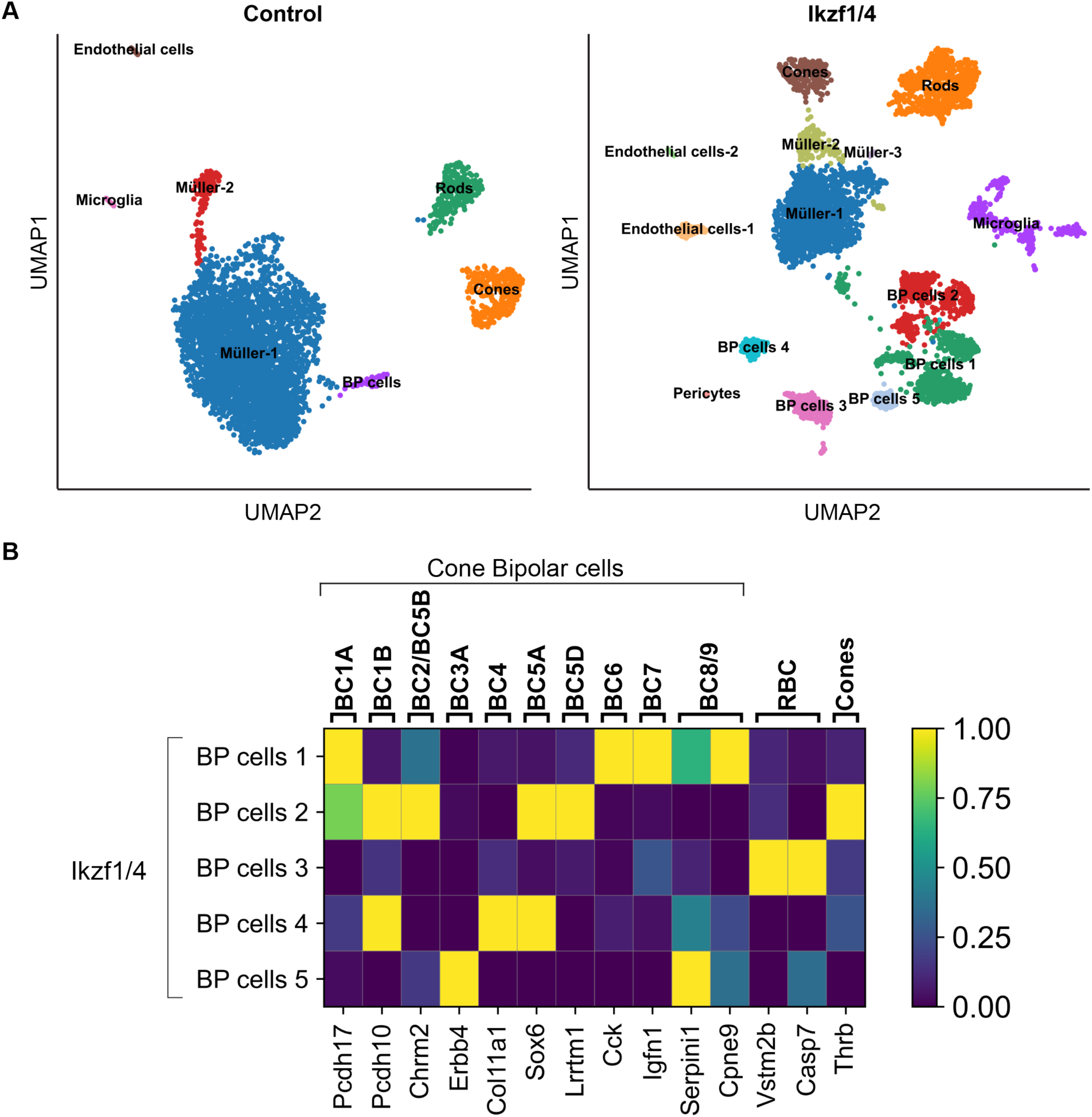
scRNA-seq identifies reprogrammed cells as bipolar-like cells. **(A)** Control UMAP (left) and Ikzf1/4 UMAP (right). BP: Bipolar. **(B)** Expression matrix of bipolar subtypes (top label) markers (bottom label) and the cone marker Thrb for Ikzf1/4 Bipolar (BP) clusters based on Shekhar et al. (Shekhar et al., 2016). Yellow denotes high expression and blue no expression. RBC: Rod bipolar cell.

To follow up on the idea that some Ikzf1/4 bipolar clusters might contain cells with both cone and bipolar transcriptional identity, we performed SCENIC analysis (Aibar et al., 2017; Van de Sande et al., 2020), which allows to investigate broad gene regulatory networks, or ‘regulons’, active within different cell populations. We found that, in addition to bipolar regulons (Fig. 5B’’), cone regulons are active in Ikzf1/4 reprogrammed cell clusters 1-7 (Fig. 5B’). Importantly, we found that the Rxrg regulon is active in Ikzf1/4 bipolar clusters (Fig. 5B’), consistent with our immunostaining data. Since we could not compare this population to endogenous bipolar cells, as they were not enriched in our control condition (Fig. 5A-A’’), we took advantage of a P14 retina scRNA-seq dataset previously published (Clark et al., 2019). As expected, bipolar cells from this dataset did not show activity of Thrb or Rxrg regulons, and showed low activity of other cone regulons (Fig. 5C’), supporting the interpretation that the Ikzf1/4 reprogrammed population represents a distinct cell type. We hence labelled these clusters as Bip/Co. Interestingly, we could not detect all MG regulons in our Ikzf1/4 dataset (Fig. 5B’’), suggesting that MG transcriptional programs are altered in this condition. Of note, the Rxrg regulon was not detected in both control datasets (Fig. 5A’, C’) potentially due to the low number of cones present and/or the low expression of Rxrg targets within this small population. Altogether, these data demonstrate that Ikzf1/4 expression in MG induces reprogramming to bipolar/cone cells.

**Figure 5:**
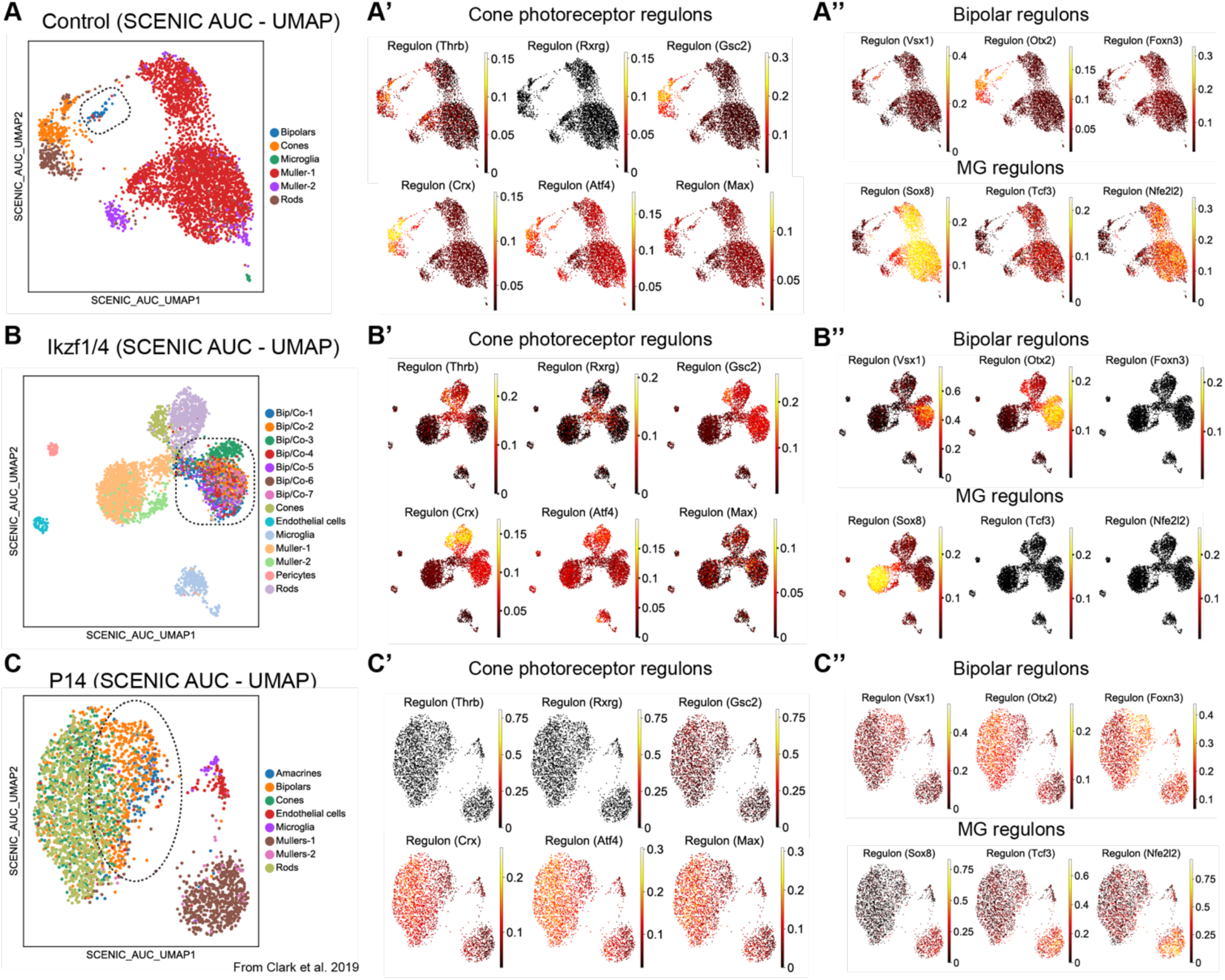
Cone and bipolar regulons are active in Ikzf1/4 reprogrammed population. **(A-C’’)** SCENIC analysis of control (A-A’’), Ikzf1/4 (B-B’’), and (Clark et al., 2019) data of P14 retinas (C-C’’) for cone photoreceptor, bipolar and MG regulons. Left: UMAP for each dataset with bipolar (or Bip/co) clusters circled. Top regulons per cluster were found by calculating the regulon specificity scores for each cluster. Shown are the top 5 enriched regulons per cluster for the P14 dataset, and Rxrg and Thrb regulon to highlight cone regulons active in the Bip/Co population. Ikzf1/4 Bip/Co have active cone (B’) and bipolar (B’’) regulons, which are not detected or present in low levels in control bipolar datasets (A’’, C’’). Scales represent AUC values (black indicates no regulon activity; Yellow represents high regulon activity).

### Ikzf1/4 reprogram mouse embryonic fibroblasts to induced neurons by remodelling chromatin and inducing a neuronal gene expression program

To investigate whether Ikzf1 and Ikzf4 have general neuronal reprogramming abilities when expressed in non-neural cells, we asked if they could convert MEFs to iNs, as previously reported for Ascl1 (Vierbuchen et al., 2010). Using a doxycycline-inducible lentiviral expression system (see methods; Fig. S6A), we expressed Ikzf1 and Ikzf4, together with Brn2 and Myt1l (BM/Ikzf1/4), and used Brn2/Ascl1/Myt1l (BAM) as positive control or Brn2/Myt1l (BM) as negative control. After 14 days, we found many Tau+ neuron-like cells in the BAM condition, as expected, and we also found Tau+ neuron-like cells in the BM/Ikzf1/4 condition, but not in the BM control condition (Fig. S6). Interestingly, expression of BM/Ikzf1 in MEFs isolated from Tau::EGFP transgenic mice resulted in upregulation of Tau::EGFP and induced neuronal morphology changes without proliferation, consistent with direct reprogramming (Video S1). These results identify Ikzf1/4 as new factors that can promote MEF to iN reprogramming.

To gain insights on the molecular mechanisms underlying Ikzf1/4 reprogramming function, we next carried out epigenetic and transcriptomic analyses on MEFs 48 hours after doxycycline-induced expression of Brn2 and Myt1l only (BM), as control, or together with Ikzf1/4 (BM/Ikzf1/4) (Fig. S6A). As classical reprogramming factors generally modify the chromatin landscape, we first investigated chromatin accessibility by ATAC-seq. We found that BM/Ikzf1/4 expression leads to a general increase in chromatin accessibility compared to the BM control condition, with open chromatin in clusters 2 and 4 forming about two thirds of all significantly altered peaks (Fig. 6A). Peak clusters 1 and 3 represent closed chromatin in BM/Ikzf1/4 compared to BM, with cluster 1 peaks showing a partial reduction in signal, and cluster 3 peaks showing almost complete loss of signal (Fig. 6A). To identify genes associated with closed and opened chromatin regions, we focused our analysis +/- 2kb from transcription start sites (TSS). Interestingly, we found that many chromatin regions that become closed after expression of Ikzf1/4 are in the cis-regulatory region of fibroblast genes, such as *Tead2*, *Ednra*, *Ogn*, *Fibin* and *Matn4*, which correspond to *transmembrane transport* gene ontology (GO) terms (Fig. 6B). In contrast, many chromatin regions that become opened after expression of Ikzf1/4 are associated with neuronal genes, such as *Gabra4*, *Gabrg1*, *Sox2*, *Neurod2*, *Pou4f1* and *Lhx5*, and are classified as general *nervous system* GO terms (Fig. 6C). Of note, numerous olfactory receptor genes were also associated with opened peaks (Table S2). These data demonstrate that expression of Ikzf1/4 in MEFs promotes chromatin accessibility of neuronal genes while reducing accessibility of fibroblast genes.

**Figure 6:**
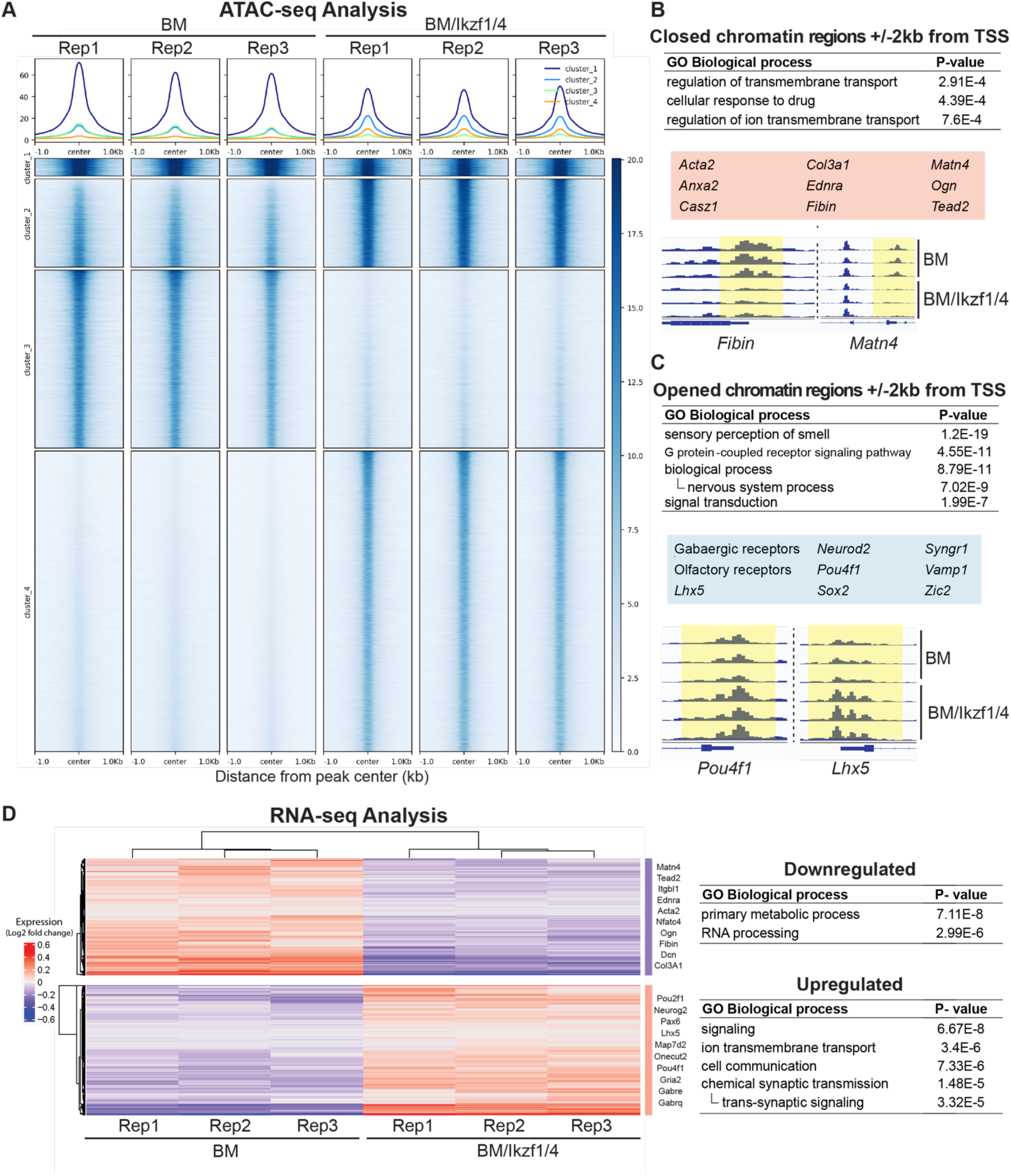
Ikzf1/4 increase chromatin accessibility and activate a neuronal gene expression program in MEFs. **(A)** ATAC signal aligned at peak center of all significantly (p<0.05) enriched or depleted ATAC peaks in BM/Ikzf1/4 (right) compared to BM (left). Clusters 1 and 3 show peaks with decreased signal in BM/Ikzf1/4 (closing of chromatin), and clusters 2 and 4 show peaks with increased signal in BM/Ikzf1/4 (opening of chromatin). Top graphs show mean signal for each cluster. Scale represents peak coverage with no coverage in white and maximum coverage in dark blue. Columns represent replicates (Rep) per infection condition. **(B, C)** Closed (B) or opened (C) chromatin regions located +/- 2kb from the transcription start site (TSS) represented as a table of GO term classification. Genes were classified using GREAT algorithm. Examples of genes associated with closed or opened chromatin regions are shown in red (B) and blue (C) boxed areas, respectively. See Table S2 for complete list of genes. TSS genomic tracks for *Fibin* and *Matn4* (B) or *Pou4f1* and *Lhx5 (C)* are shown in the bottom panels. **(D)** Heatmap of log2 expression fold change and GO term classification of the most significantly upregulated and downregulated genes in MEFs 48 hours after BM (left) or BM/Ikzf1/4 (right) induced expression. Parameters used for scoring significant genes were Log2FC>0.25 and p-value<0.05. High expression is denoted by red whereas low expression is denoted by blue. Each replicate is represented as a column (n=3). Examples of downregulated fibroblast genes and upregulated neuronal genes in BM/Ikzf1/4 compared to BM are listed on the right side of the heatmap. GOrilla classification of GO terms are represented as a table on the right.

We next assessed whether this change in chromatin architecture resulted in altered gene expression by RNA-seq. We found that expression of BM/Ikzf1/4, compared to BM, leads to significant upregulation of several neuronal genes like *Neurog2*, *Pax6*, *Lhx5*, *Pou4f1*, and some GABA and glutamate receptors (Fig. 6D; Table S3), suggesting that both gabaergic and glutamatergic neurons are generated. Notably, Ikzf1/4 also upregulates expression of *Sall3*, *Pou2f1* and *Onecut2*, three genes that were previously linked to cone photoreceptor development (de Melo et al., 2011; Javed et al., 2020; Sapkota et al., 2014), as well as *Lhx4* and *Isl1*, which are involved in bipolar cell specification (Table S3) (Dong et al., 2020; Elshatory et al., 2007). *Ascl1* expression was not upregulated in the BM/Ikzf1/4 condition, suggesting that Ikzf1/4 does not require Ascl1 expression to induce neuronal reprogramming. Upregulated genes were associated with GO terms like *cell-cell signaling, ion transmembrane transport,* and *chemical synaptic transmission* (Fig. 6D). Conversely, we observed downregulation of several fibroblast genes like *Matn4*, *Tead2*, *Nfatc4*, and *Fibin* (Fig. 6D, Table S3). These genes belong to GO terms associated with various *metabolic processes* (Fig. 6D). Of note, olfactory receptors that were found to have opened chromatin after Ikzf1/4 expression did not generally show an increase in transcript levels, suggesting they are not involved in reprogramming. Overall, approximately 17% of genes associated with BM/Ikzf1/4-enriched ATAC peaks show increased transcript levels, including the neuronal genes *Sall3*, *Gabra4*, *Lhx5*, *Pou4f1*, and *Zic2* (Fig. S7, Table S4). A similar percentage of genes associated with reduced ATAC peaks show decreased transcript levels, including MEF genes *Ednra*, *Fibin, Matn4*, and *Tead2* (Fig. S7, Table S4). Together, these results demonstrate that Ikzf1/4 quickly inhibit fibroblast and activate neuronal transcriptional programs in MEFs.

## DISCUSSION

Vertebrate temporal identity factors control competence of neural progenitors during development, but here we show that they can additionally reprogram the identity of fully differentiated cells, linking the concepts of temporal identity transitions in progenitors with that of somatic cell states. Specifically, we show that the early temporal identity factor Ikzf1 is a potent reprogramming factor capable of converting adult retinal glia to bipolar/cone-like cells when co-expressed with Ikzf4 (Fig. 7A), and MEFs to iNs when co-expressed with Ikzf4, Brn2 and Myt1l (Fig. 7B). This work uncovers reprogramming properties for temporal identity factors and a previously unappreciated reprogramming pathway that could lead to new therapeutic opportunities.

**Figure 7:**
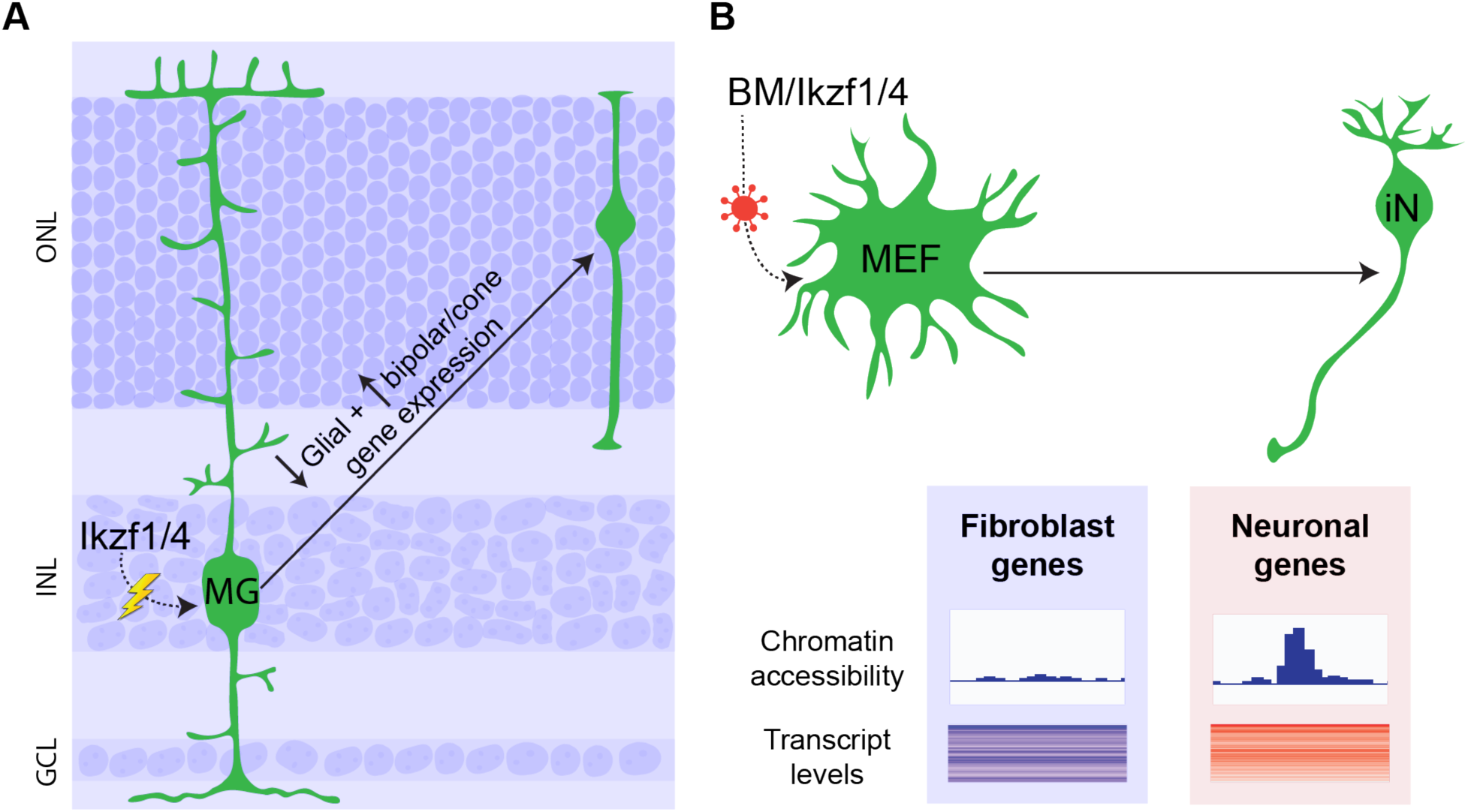
Summary of findings. **(A)** Ikzf1/4 expression in MG through electroporation, represented by lightning bolt and dotted line, induces reprogramming (arrow) into cone-like cells located in the photoreceptor layer. The reprogrammed cells turn off glial gene expression and start expressing both cone and bipolar cell genes. **(B)** BM/Ikzf1/4 expression in MEFs through lentiviral vectors, represented by dotted line and red virus cartoon, reprograms (arrow) these cells to iNs by decreasing accessibility and expression of fibroblast genes (represented in blue), and increasing accessibility and expression of neuronal genes (represented in red). iNs: induced neurons.

### Eliciting the reprogramming potential of temporal identity factors

The idea brought forward here, that temporal identity factors may not only control progenitor identity but also mediate reprogramming of differentiated cells, has previously been explored in *Drosophila*. Pearson and Doe (Pearson and Doe, 2003) showed that expression of *hb* in late-born neurons cannot convert them to early-born neuronal cell types, whereas more recent work showed that expression of hb in neuroblasts past the fifth division is also insufficient to confer early competence, probably due to chromatin remodelling that renders older neuroblasts and differentiated cells unresponsive to *hb* expression (Kohwi et al., 2013). Consistently, we show here that the mammalian homolog of *hb*, Ikzf1, cannot reprogram MG when expressed on its own. However, we find that co-expression of Ikzf1 with Ikzf4 elicits a reprogramming response in MG. Ikzf1 and Ikzf4 are known to physically interact (Honma et al., 1999), and this interaction may potentiate their reprogramming ability. As Ikzf4 is expressed in the developing mouse retina (Clark et al., 2019; Elliott et al., 2008), these results also raise the possibility that Ikzf4 might cooperate with Ikzf1 to control temporal identity of retinal progenitor cells.

We screened several other factors that might promote MG reprogramming, including temporal identity factors like Casz1v2 and Pou2f1, but we find that no other combination leads to cell identity conversion. It is possible that appropriate potentiating factors were missing in these experiments, limiting our ability to elicit the reprogramming potential of these temporal factors. For instance, the late temporal identity factor Casz1 strongly promotes rod photoreceptor fate during development (Mattar et al., 2015), raising the possibility that co-expression of rod differentiation factors like Otx2, Crx, or Nrl may be necessary to unravel Casz1 reprogramming ability. Other factors may also allow Ikzf1 reprogramming towards different early-born cell types, such as retinal ganglion cells, horizontal cells, or amacrine cells, which are ectopically-produced by late retinal progenitors upon expression of Ikzf1 (Elliott et al., 2008). Co-expression of factors involved in cone differentiation, such as Sall3 (de Melo et al., 2011) or Crx (Chen et al., 1997; Furukawa et al., 1997), may also be necessary to engage Ikzf1/4 reprogrammed cells into a complete cone differentiation pathway. Our data indicate that Ikzf1/4 expression in MG confers at least some cone reprogramming competence. This is an exciting achievement, as these findings could represent a foundation for future work aimed at finding conditions to reprogram MG into functional cones to develop therapies for cone photoreceptor degenerative diseases.

Interestingly, Fezf2, a transcription factor regulating cortical progenitor output, was shown to reprogram late-born neurons to an early-born neuronal cell type (De la Rossa et al., 2013; Rouaux and Arlotta, 2013). Although Fezf2 reprogramming was limited to young neurons and changed neuronal subtype rather than cell type identity, the ability of this transcription factor to alter the identity of differentiated cells parallels our work. It will also be interesting to determine whether cell identity reprogramming can be achieved in the neocortex by expression of temporal identity factors.

### Mechanisms of Ikzf1/4-mediated direct neuronal reprogramming

Many lines of evidence reported here support that Ikzf1/4 function as direct reprogramming factors, without inducing a pluripotent or proliferative state. First, ex vivo EdU experiments show that expression of Ikzf1/4 in MG does not stimulate re-entry into the cell cycle. Second, in vivo expression of Ikzf1/4 in MG leads to isolated reprogrammed cells in the photoreceptor layer, but if proliferation occurred we would expect reprogrammed cells to agglomerate in small clusters. Third, we do not find MG positive for progenitor markers by immunostaining, nor clusters of neurogenic progenitors in the scRNA-seq analyses. Fourth, timelapse video recordings show that MEFs do not undergo division before converting to iNs. Taken together, these data support a mechanism of direct conversion by which MG and MEFs turn off glia or fibroblast genes, respectively, and turn on neuronal genes following expression of Ikzf1/4 (Fig. 7). Although this contrasts with teleost fish MG-mediated regeneration, where MG de-differentiate into a proliferating progenitor state before re-differentiating into retinal neurons (Goldman, 2014), it is concordant with previous work in mammals showing MG reprogramming to neuron-like cells without proliferation (Hoang et al., 2020; Jorstad et al., 2017). Of note, temporal identity factors change competence of neural progenitors without altering their proliferative capacity (Elliott et al., 2008; Javed et al., 2020; Mattar et al., 2015). Our data provide evidence that this property is maintained when temporal factors are expressed in differentiated cells.

We also report that Ikzf1/4 expression in MEFs leads to widespread chromatin reorganization, as observed with most classical reprogramming factors studied to date (Dall’Agnese et al., 2019; Koche et al., 2011; Wapinski et al., 2017; Wapinski et al., 2013), providing a potential molecular mechanism for Ikzf1/4-mediated reprogramming. Reprogramming factors usually have pioneer activity, binding and opening chromatin regions that are normally not accessible to transcription factors (Iwafuchi-Doi and Zaret, 2016). While it remains unclear whether Ikzf1/4 function as pioneer factors, the widespread changes in chromatin accessibility observed within 48 hours after expression in MEFs is consistent with this possibility. In such context, Ikzf1 might play a critical role, as it is known to interact with chromatin remodeling complexes such as Mi-2/NuRD and SWI/SNF (Kim et al., 1999; O’Neill et al., 2000). The NuRD complex was found to oppose reprogramming to pluripotency (Mor et al., 2018; Rais et al., 2013), supporting the notion that Ikzf1/4 promote direct transdifferentiation without inducing a pluripotent state. We recently reported that the late temporal identity factor Casz1 also interacts with the NuRD complex and requires polycomb activity to control the neurogenesis to gliogenesis transition in retinal progenitors (Mattar et al., 2021), suggesting that altering chromatin state may be a common theme for vertebrate temporal identity factors. Conversely, in *Drosophila*, secreted spatial patterning factors, rather than temporal factors, control chromatin accessibility states in different neural progenitors. This allows the same temporal identity factor to bind different transcriptional targets and alter fate output differently depending on the location of the neuroblast in which it is expressed (Sen et al., 2019). These results suggest that temporal factors use different mechanisms to alter progenitor competence in vertebrates and invertebrates. The chromatin-modifying ability of vertebrate temporal identity factors might explain why Ikzf1/4 can elicit direct neuronal reprogramming, whereas *hb* cannot (Kohwi et al., 2013). Importantly, our data indicate that Ikzf1/4 expression not only alters chromatin accessibility, but that this translates into activation of a neuronal transcriptional program and repression of fibroblast gene expression.

Ikaros family members, including Ikzf1 and Ikzf4, are well known for their role in hematopoiesis (Heizmann et al., 2017; John and Ward, 2011). Yet, we find that most upregulated genes after Ikzf1/4 expression in MEFs are associated with neuronal and not hematopoiesis processes. It is possible that Brn2 and Myt1l direct Ikzf1/4 towards neuronal rather than hematopoiesis targets. Indeed, Myt1l has been shown to suppress non-neuronal gene expression (Mall et al., 2017). Interestingly, we also find that Ikzf1/4 control chromatin conformation at olfactory receptor loci. Although this does not result in altered expression of olfactory receptor genes in MEFs, it suggests that Ikzf1/4 could be involved in olfactory neuron specification by regulating chromatin accessibility of olfactory receptor promoters. Ikzf1 binding sites were previously identified at promoter regions of olfactory receptors (Lane et al., 2001; Plessy et al., 2012), but its role in olfactory neuron production is unknown.

Reprogramming properties were also recently reported for Onecut factors in MEFs (van der Raadt et al., 2019). Interestingly, we observe here that Ikzf1/4 expression in MEFs induces upregulation of *Onecut2* within 48 hours, suggesting that these factors might act through a common program. In contrast, we find that Ikzf1/4 expression does not upregulate *Ascl1* in MEFs, supporting an Ascl1-independent reprogramming mechanism. To date, most studies in fibroblasts take advantage of Ascl1 to promote reprogramming, and use co-expression of various neural transcription factors to induce differentiation into specific neuronal subtypes for disease modeling and cell transplantation (Chen et al., 2019). Together with recent results of large-scale screens (Liu et al., 2018; Tsunemoto et al., 2018), the identification of Ikzf1/4 as Ascl1-independent reprogramming factors described here should provide a foundation for future work aimed at increasing the diversity of neuronal cell types that can be generated by direct reprogramming.

### Temporal identity factor-mediated reprogramming in the CNS

Many temporal identity factors, including Ikzf1, remain expressed in differentiated cells (Elliott et al., 2008; Javed et al., 2020; Mattar et al., 2015), but the functional implication of this is still unclear. The identification of broad reprogramming abilities of Ikzf1 reported here suggests that deregulation of Ikzf1 expression in differentiated cells may lead to abnormal cell states. Whether this occurs endogenously in the mammalian CNS or in various disease conditions remains to be investigated.

Temporal identity factor-mediated neuronal reprogramming is a potentially broadly applicable technique that may elicit a regenerative response from various CNS glia. As Ikzf1 also regulates early temporal identity in cortical progenitors and is sufficient to promote early-born neuron production when expressed in late progenitors (Alsio et al., 2013), it is tempting to speculate that it might act as a reprogramming factor in cortical astrocytes. Still, it is important to note that MG have a similar transcriptome to late retinal progenitors (Blackshaw et al., 2004; Jadhav et al., 2009; Roesch et al., 2008), which might make them more prone to temporal identity factor-mediated reprogramming than other cell types. But reactive astrocytes also show some progenitor-like properties and gene expression profile (Gotz et al., 2015), similar to retinal MG (Hoang et al., 2020), suggesting they may be susceptible to reprogramming with temporal identity factors.

Together with previously published results, the findings reported here support a conceptual similarity between temporal and cell identity reprogramming, identifying a broadly applicable method for neuronal reprogramming with potential implications for nervous system development and regeneration.

## ACKNOWLEDGEMENTS

We would like to thank Drs. Seth Blackshaw and Brian Clark for comments on the manuscript, and Javier Di Noia for providing us an Apobec2b construct. We thank the IRCM core facilities and personnel for technical assistance, in particular Jessica Barthe, Marie-Claude Lavallée, Julie Mercier, and Caroline Dubé for animal care, Dominic Filion for miscroscopy, Julie Lord for flow cytometry, and Odile Neyret for next generation sequencing. We would also like to thank Christine Jolicoeur for technical assistance and the entire Cayouette team for feedback and suggestions throughout the realization of this work. C.B.P and A.J. received scholarships from the Fonds de recherche du Québec – Santé, and C.B.P. from the Canadian Institutes of Health Research. This project was funded by grants from the Canadian Institutes of Health Research (FDN-159936), Fighting Blindness Canada, and the Krembil Foundation to M.C.. M.C. is an Emeritus Scholar from Fonds de recherche du Québec – Santé, and holds the Gaëtane and Rolland Pillenière Chair in Retinal Biology from the Montreal Clinical Research Institute Foundation.

## AUTHOR CONTRIBUTIONS

Conceptualization: C.B.P., M.F., P.M., M.C. ; Investigation and analysis: C.B.P., A.J., M.F.; Resources: M.C.; Writing - original draft: C.B.P.; Writing - reviewing and editing: C.B.P., P.M., M.C.; Supervision: M.C.; Funding acquisition: M.C.

## MATERIALS AND METHODS

### Animals

Animal work was performed in accordance with the Canadian Council on Animal Care and IRCM guidelines. Tg(Slc1a3-cre/ERT)1Nat/J mice (Glast-CreER, stock 012586), B6.129X1-*Gt(ROSA)26Sor^tm1(EYFP)Cos^*/J reporter mice (R26R-EYFP, stock 006148), C3H/HeJ mice (Pde6b^RD1/RD1^, stock 000659), B6.Cg- *Gt(ROSA)26Sor^tm1(rtTA*M2)Jae^*/J mice (R26-M2rtTA, stock 006965), Mapt^tm1(EGFP)Klt^/J (Tau- EGFP, stock 004779), and C57BL/6J (stock 000664) were all obtained from The Jackson Laboratory. Glast-CreER mice were always heterozygous, R26R-EYFP mice were fl/+ for mouse line validation and ex vivo work and fl/+ or fl/fl for in vivo work.

### DNA constructs

pCALL2 (https://health.uconn.edu/mouse-genome-modification/resources/conditional-knock-outexpression-vectors/) was digested with ClaI and SphI to remove LacZ and insert mCherry (amplified from MSCV-mCherry) in the loxP cassette. IRES-EGFP was removed with SmaI and NotI digestions to generate a loxP-mCherry-loxP-Multiple Cloning Site (MCS) construct, which we named pCALM to distinguish from the original pCALL2. A Gateway cassette was added within the MCS to allow insertions of some coding sequences with Gateway Cloning System (Thermo Fisher), while others were inserted directly in the MCS by restriction digestions or with In-Fusion cloning (Clontech). Empty pCALM-MCS was used as control for retinal work. Construct was tested in HEK 293T (ATCC) cells by jetPRIME (Polyplus) transfection alone or with pCIG-Cre following instructions from manufacturer.

psPAX2 (Addgene plasmid # 12260; http://n2t.net/addgene:12260; RRID:Addgene_12260) and pMD.2G (Addgene plasmid # 12259; http://n2t.net/addgene:12259; RRID:Addgene_12259) were provided by Dr. Didier Trono. TeT-O-FUW-Ascl1 (Addgene plasmid # 27150; http://n2t.net/addgene:27150; RRID:Addgene_27150), TeT-O-FUW-Brn2 (Addgene plasmid # 27151; http://n2t.net/addgene:27151; RRID:Addgene_27151) and TeT-O-FUW-Myt1L (Addgene plasmid # 27152; http://n2t.net/addgene:27152; RRID:Addgene_27152) were provided by Dr. Marius Wernig (Vierbuchen et al., 2010), whereas TeT-O-FUW-Ikzf1 and TeT-O-FUW-Ikzf4 were generated by cloning the Ikzf1 and Ikzf4 coding sequences into TeT-O-FUW (obtained by removing Brn2 from TeT-O-FUW-Brn2) by standard techniques. These plasmids were transformed in Stbl2 competent cells (Thermo Fisher) to prevent recombination with bacterial DNA due to the repetitive sequences of the lentiviral backbones.

### Glast-CreER;R26R-EYFP mouse line validation

P30-40 Glast-CreER;R26R-EYFP mice received tamoxifen diluted in canola oil (Toronto Research Chemicals or Cedarlane Labs) by intraperitoneal injections for four consecutive days at 90µg/gram of body weight. Mice were euthanized by CO_2_ at three to four days after the last tamoxifen injection.

### Ex vivo reprogramming

Eyes from P0-1 Glast-CreER;R26R-EYFP mice were collected in DPBS (Gibco) under sterile conditions. Retinas were electroporated with single or combined plasmids, with pCALM-MCS as control. Vectors (1µl at 3µg/µl) were injected in the sub-retinal space and a current (50millisec duration, 950 millisec interval, 40-50 volts, unipolar electrodes; BTX ECM 830) was applied over the eye with the positive electrode facing the cornea. Retinas were then dissected in DPBS and placed on a culture insert (Millicell) in a 6-well plate (Falcon) containing 1.3ml of equilibrated medium: DMEM (supplemented with glutaMAX; Gibco) with 10% FBS (Sigma) and 1x Pen/Strep (Gibco). At DIV12, hydroxy-tamoxifen (Cayman Chemical Co.) and EGF (PreproTech) were added to the culture medium at a final concentration of 5µM and 100ng/ml, respectively. Two to three days later (DIV14/15), the solution was removed and replaced with fresh culture medium (DMEM/10%FBS/Pen/Strep) after rinsing the well with DPBS. In some experiments, 2’-Deoxy-5-ethynyluridine (EdU, Abcam) was added to the culture medium at a concentration of 10µg/ml for three days. At DIV26, the solution was removed and replaced with 1ml of 4% paraformaldehyde (PFA; Electron microscopy sciences) in PBS for five minutes at room temperature. 1ml of 4% PFA was then added over the culture insert and incubated for another 5-minute at room temperature. Explants were quickly washed with PBS and immersed in 20% sucrose in PBS at 4°C for 2 two to five hours before being detached from the insert with curved forceps and frozen in a 20% sucrose:OCT (1:1, Sakura).

### In vivo reprogramming

Glast-CreER;R26R-EYFP P0-2 mice were anesthetized on ice, injected sub-retinally with 1µl of DNA vectors (3µg/µl; pCALM-MCS or pCALM-Ikzf1 + pCALM-Ikzf4) in one eye and subjected to an electrical current as previously described (de Melo and Blackshaw, 2011). In some experiments, animals were injected intraperitoneally with EdU at 50µg/g of body weight daily from P3-7. Animals were injected intraperitoneally or gavaged daily with 90µg of tamoxifen/gram of body weight for three consecutive days, starting any day between P21 and P35. Animals were euthanized by CO_2_ three to five weeks post-tamoxifen, as specified. Eyes were collected and the retinas dissociated for scRNA-seq (see cell sorting section) or fixed and processed for immunofluorescence (see below).

### Primary mouse embryonic fibroblast (MEF) culture

Homozygous R26-M2rtTA male mice were crossed with wildtype C57BL/6J or Tau-EGFP females. Plugged females were sacrificed at E13.5 and embryos extracted. Primary MEF culture was performed as previously described (Jozefczuk et al., 2012) with some modifications: 0.25% Trypsin-EDTA (Thermo Fisher) was used for dissociation, and dissociated cells were left for 15 minutes at room temperature to let cell debris and remaining tissue sink to the bottom of the tube, the supernatant was collected and centrifuged. MEFs were cultured in MEF medium: DMEM, 10% Heat-inactivated Cosmic Calf Serum (Cytiva), 1% MEM non-essential amino acids solution (100X stock; Thermo Fisher), 1nM Sodium-Pyruvate (Thermo Fisher), 100U/ml Pen/Strep (Thermo Fisher), 0.114mM 2-Mercaptoethanol (Sigma). When cells reached 90-100% confluency, they were either passaged or frozen in liquid nitrogen. Cells were passaged at least three times before lentiviral vector infection.

### Lentiviral vectors production

293FT cells (Thermo Fisher) were plated onto 10cm dishes and transfected at 70% confluency. For transfection, Polyethylenimine (PEI 25K, Polysciences) was added to 1ml of DMEM at a final concentration of 45ng/ul with 5µg of psPAX2, 10µg of pMD.2G, TeT-O-FUW-Ascl1, TeT-O-FUW-Brn2, TeT-O-FUW-Myt1L, TeT-O-FUW-Ikzf1 or TeT-O-FUW-Ikzf4, incubated for 15 minutes at room temperature, and the solution was added drop-wise to the cells. Six hours later, the culture medium was replaced with fresh DMEM supplemented with 5% BSA (Sigma Aldrich). Lentiviral collection and spindown was performed at 24 hours and 48 hours after initial change of medium with Lenti-X-concentrator (Clontech) according to manufacturer protocol. Viral pellet was resuspended in DMEM, aliquoted and stored at -80°C. When ready to use, lentiviral vector aliquots were thawed rapidly at room temperature, centrifuged at 3000g for three minutes and the supernatant used for infection.

### Lentiviral infection of MEFs

MEFs were seeded in a 6-well plate at 200 000 cells/well. Infection was carried out about 12 hours post-plating if the cells had reached a confluency of at least 50%. Prior to infection, cells were incubated at 37°C for 30-45 minutes in MEF medium (described in MEF culture section) containing 8µg/ml of Polybrene (Santa Cruz Biotechnology). Ascl1, Brn2, Myt1l, Ikzf1, and Ikzf4 lentiviral vectors were added to each corresponding condition to obtain a multiplicity of infection (MOI) of 2. Medium containing lentiviral vectors was discarded and replaced with fresh MEF medium 12-18 hours after infection. 24 hours later, cells for immunostaining were trypsinized and replated on acid-washed glass coverslips coated with 0.1% bovine gelatin (Sigma) in 24-well plates while cells for ATAC-seq and RNA-seq were maintained in 6-well plates. MEF medium containing 10µM of the caspase-inhibitor quinoline-val-asp-difluorophenoxymethylketone (QVD-OPH; MedChem Express) was added to the cells at this time.

### MEF reprogramming assay

MEF reprogramming assay followed a previously published protocol (Vierbuchen et al., 2010) with some modifications. 24 hours after plating, the culture medium was changed to MEF medium containing 10µM QVD-OPH and 2µg/ml doxycycline (Sigma), corresponding to D0 of the assay. 48 hours later, MEF medium was replaced with reprogramming medium for the rest of the assay, with changes every two days: DMEM/F12 (Gibco) with 1/50 B27-Supplement without vitamin A (50X stock; Thermo Fisher), 1/100 N2-Supplement (homemade, as previously reported by (Vierbuchen et al., 2010)), 2µg/ml doxycycline, 10µM QVD-OPH, 5ng/ml BDNF, 10ng/ml CNTF, and 10ng/ml NT-3 (PeproTech). At D14, cells were washed with PBS and incubated for 15 minutes in 4% PFA/PBS solution at room temperature. For time-lapse recording, M2rtTA;Tau-EGFP MEFs were cultured in an Incucyte Live-Cell Analysis incubator (Sartorius) for 12 days, starting with doxycycline addition to the culture medium. Pictures were taken every 8 hours.

### Eye fixation and cryosectionning

Eyes were collected and fixed five minutes at room temperature for reprogramming work or three to six hours at 4°C for mouse line validation in 4% PFA/PBS, washed with PBS, and immersed in 20% sucrose in PBS for four hours to overnight at 4°C, then frozen in 20% sucrose:OCT (1:1) for cryosectionning. Retinal explants and eyes were sectioned with a cryostat (Leica) at 16 µm for mouse line validation or 25µm for other assays.

### Immunostaining

Immunostaining for retinal explant and eye sections were performed as previously described (Javed et al., 2020). Slides stained for CyclinD3 underwent a two hour antigen retrieval in sodium citrate buffer at 54°C before blocking incubation. See Method Table S1 for primary antibody list. When specified, slides were processed for EdU Click-It reaction following the manufacturer protocol (Abcam; modified to use half of recommended AlexaFluor-647).

MEFs were incubated for 30 minutes in blocking solution: 3% BSA and 0.5% Triton-X in PBS, and then incubated at room temperature for two and a half hours in blocking solution with primary antibodies (See Methods Table 1). Cells were then washed 3x in PBS and incubated in blocking solution with secondary antibodies (1/1000) and Hoechst (1/10000) at room temperature for one to two hours. Cells were then washed 3x in PBS and the coverslips mounted on a microscopy slide with Mowiol for analysis.

### Microscopy and cell counts

For Glast-CreER;R26R-EYFP line validation, three to four images per animal and each marker were randomly acquired using a 40x objective with a Leica DM6000 microscope and analyzed on Volocity™ software (Perkin Elmer) to quantify the number of YFP+ cells co-labelling with cell-type specific markers. For reprogramming assays, images were acquired using a 20x or 63x objective on an SP8 confocal microscope (Leica), analyzed on Volocity™ software, and processed on Fiji™ (ImageJ), and Adobe™ Illustrator (Adobe). Images in figures are z-projections or single planes, as specified in figure legends, for optimal representation of cell morphologies. Electroporated (mCherry+) regions with normal retinal morphology were selected for quantification. For cell counts, YFP+ mCherry+ cells were analyzed. MEFs were imaged using a Leica DM6 microscope.

### Fluorescent-activated cell sorting

Glast-CreER;R26R-EYFP retinas were isolated and mCherry regions were dissected out using a mini scalpel under a fluorescent microscope (Leica MZ16FA). Retina pieces were dissociated for 16 minutes in papain (82.5U; Worthington) at 37°C. The reaction was stopped by rinsing the retinas once with Lo-Ovo (DPBS, 1.5% BSA, 1.5% Trypsin inhibitor (Roche Diagnostics), pH 7.4) solution with DNase (Worthington Biochemical; 0.3U/ul), and then re-suspended in Lo-Ovo with DNase solution for gentle trituration by pipetting up and down slowly with a P1000 pipet (Gilson). The cell suspension was then passed through a 70µm filter and Hoechst 33258 (Invitrogen) was added to the solution to label dying cells. Fluorescent cells were sorted with a FACSAria III Cell Sorter (BD Biosciences) with a 100µm nozzle. Viable YFP+ cells (single cells) were collected in PBS with 0.15% BSA (Millipore). YFP+ cells from mCherry+ regions of 7 retinas (n=7) were pooled for the Ikzf1/4 condition. As control, YFP+ cells from both mCherry+ (n=4 retinas) or mCherry- (n=5 retinas) regions were pooled. Cells were then spun at 300g for 10 minutes at 4°C and re-suspended in 0.15% BSA/PBS.

### scRNA-sequencing

Cells were loaded on a 10xGenomics Single Cell 3’ chip and processed according to manufacturer V3 pipeline 3.1.0 kit. cDNA libraries we sequenced with Illumina NovaSeq 6000 at an estimated 20000 reads per cell. A total of 4207 cells were sequenced for the control sample and 4608 cells for the Ikzf1/4 sample. Data can be found on GEO #GSE169519.

### scRNA-sequencing analysis

scRNA-seq fastq files were processed with Cellranger version 4.0 to generate counts with default parameters and reference index provided by 10x Genomics (refdata-cellranger-mm10-2020-A). Loom files were generated with Velocyto with run10x function on the cellranger folder and repeat mask for mm10 genome from UCSC genome browser (La Manno et al., 2018). Scanpy was used to analyze loom files and generate UMAP clusters for both conditions. Scanpy Matrix plot function was used to assess cell type specific markers in each UMAP cluster. Marker genes were chosen from previously published scRNA-seq datasets of mouse retina and sorted Bipolar cells (Clark et al. 2019, Shekhar et al. 2016). Code can be found on github (https://github.com/awaisj14/BipCo).

Single cell regulatory network inference and clustering (SCENIC) analysis was conducted on the same loom files generated from Velocyto. Default parameters for mm10 RcisTarget database were used according to the SCENIC vignette (https://github.com/aertslab/SCENIC) (Aibar et al., 2017; Van de Sande et al., 2020). Cell type clustering and regulon AUC UMAPs were generated with Scanpy scatter plot. Code can be found on github (https://github.com/awaisj14/BipCo).

### RNA-isolation and sequencing

RNA extraction from MEF cultures was performed using the RNeasy Plus Micro Kit (Qiagen). At 48 hours after doxycycline addition, cells were washed briefly with room temperature DPBS and cells collected by scrapping the well surface with a 1ml pipette tip containing 450µl cold RLT buffer. Cell suspension was transferred in 1.5ml DNA-LoBind Eppendorf tubes. Further extraction steps were performed according to manufacturer guidelines. mRNA was isolated by Poly(A) mRNA magnetic isolation (NEBNext; E7490S), cDNA libraries were generated with KAPA RNA HyperPrep kits (Roche; 08098197702), and sequenced on the Illumina NovaSeq 6000 sequencing system. Data can be found on GEO: GSE169519.

### RNA-sequencing analysis

Salmon quant function was used to quantify effective length of transcripts and transcript per million value from RNA-seq raw fastq reads (Patro et al., 2017) and the Galaxy platform was used to perform downstream RNA-seq analysis (Afgan et al., 2016). Counts normalization and differential expression analysis was performed with DESeq2 (Love et al., 2014) and heatmaps were generated with Complex Heatmap (Gu et al., 2016). GOrilla was used to classify genes in GO classification terms (Eden et al., 2009) and REVIGO was used to summarize the most significantly enriched terms (Supek et al., 2011).

### ATAC-sequencing

48 hours after doxycycline addition, 50000 MEF nuclei were isolated as previously reported (Buenrostro et al., 2015), with modifications described in (Mayran et al., 2018). Isolated nuclei underwent transposase reaction: 2.5μl of 10x TD buffer, 10μl of water, and 12.5μl of enzyme (Illumina Nextera kit; FC-121–1031). DNA was then purified with GeneRead Purification columns (Qiagen), enriched by PCR, and purified again with GeneRead Purification columns before being sequenced with Illumina NovaSeq 6000. Data can be found on GEO: GSE169519.

### ATAC-sequencing analysis

Raw ATAC-seq fastq reads were aligned with bowtie2 on the Galaxy platform (Afgan et al., 2016; Langmead and Salzberg, 2012) and peak calling was performed with MACS2 with the following parameters: --nomodel --shift -37 --extsize 73 (Feng et al., 2012). Bamcoverage was used to generate bigwig files and Integrative Genomics Viewer (IGV) for visualization of ATAC peaks (Ramirez et al., 2016; Robinson et al., 2011). Deeptools was used to compute peaks in a matrix using computeMatrix and heatmaps were generated using plotHeatmap (Ramirez et al., 2016). GO term classification +/-2kb from TSS was performed with GREAT algorithm (McLean et al., 2010) and REVIGO was used to summarize the most significantly enriched terms (Supek et al., 2011).

### Statistics

Statistical analyses are described in figure legends. All statistical tests were performed using Prism (GraphPad) software.

## REFERENCES

Afgan, E., Baker, D., van den Beek, M., Blankenberg, D., Bouvier, D., Cech, M., Chilton, J., Clements, D., Coraor, N., Eberhard, C., et al. (2016). The Galaxy platform for accessible, reproducible and collaborative biomedical analyses: 2016 update. Nucleic acids research 44, W3–W10.

Aibar, S., Gonzalez-Blas, C.B., Moerman, T., Huynh-Thu, V.A., Imrichova, H., Hulselmans, G., Rambow, F., Marine, J.C., Geurts, P., Aerts, J., et al. (2017). SCENIC: single-cell regulatory network inference and clustering. Nature methods 14, 1083–1086.

Alsio, J.M., Tarchini, B., Cayouette, M., and Livesey, F.J. (2013). Ikaros promotes early-born neuronal fates in the cerebral cortex. Proceedings of the National Academy of Sciences of the United States of America 110, E716–725.

Alunni, A., and Bally-Cuif, L. (2016). A comparative view of regenerative neurogenesis in vertebrates. Development 143, 741–753.

Barker, R.A., Gotz, M., and Parmar, M. (2018). New approaches for brain repair-from rescue to reprogramming. Nature 557, 329–334.

Blackshaw, S., Harpavat, S., Trimarchi, J., Cai, L., Huang, H., Kuo, W.P., Weber, G., Lee, K., Fraioli, R.E., Cho, S.H., et al. (2004). Genomic analysis of mouse retinal development. PLoS biology 2, E247.

Blackshaw, S., and Sanes, J.R. (2021). Turning lead into gold: reprogramming retinal cells to cure blindness. Journal of Clinical Investigation 131, e146134.

Boudreau-Pinsonneault, C., and Cayouette, M. (2018). Cell lineage tracing in the retina: Could material transfer distort conclusions? Dev Dyn 247, 10–17.

Bringmann, A., Iandiev, I., Pannicke, T., Wurm, A., Hollborn, M., Wiedemann, P., Osborne, N.N., and Reichenbach, A. (2009). Cellular signaling and factors involved in Muller cell gliosis: neuroprotective and detrimental effects. Progress in retinal and eye research 28, 423–451.

Brody, T., and Odenwald, W.F. (2000). Programmed transformations in neuroblast gene expression during Drosophila CNS lineage development. Dev Biol 226, 34–44.

Buenrostro, J.D., Wu, B., Chang, H.Y., and Greenleaf, W.J. (2015). ATAC-seq: A Method for Assaying Chromatin Accessibility Genome-Wide. Curr Protoc Mol Biol 109, 21 29 21-29.

Chanda, S., Ang, C.E., Davila, J., Pak, C., Mall, M., Lee, Q.Y., Ahlenius, H., Jung, S.W., Sudhof, T.C., and Wernig, M. (2014). Generation of induced neuronal cells by the single reprogramming factor ASCL1. Stem cell reports 3, 282–296.

Chen, S., Wang, Q.-L., Nie, Z., Sun, H., Lennon, G., Copeland, N.G., Gilbert, D.J., Jenkins, N.A., and Zack, D.J. (1997). Crx, a Novel Otx-like Paired-Homeodomain Protein, Binds to and Transactivates Photoreceptor Cell-Specific Genes. Neuron 19, 1017–1030.

Chen, S., Zhang, J., Zhang, D., and Jiao, J. (2019). Acquisition of functional neurons by direct conversion: Switching the developmental clock directly. J Genet Genomics 46, 459–465.

Clark, B.S., Stein-O’Brien, G.L., Shiau, F., Cannon, G.H., Davis-Marcisak, E., Sherman, T., Santiago, C.P., Hoang, T.V., Rajaii, F., James-Esposito, R.E., et al. (2019). Single-Cell RNA-Seq Analysis of Retinal Development Identifies NFI Factors as Regulating Mitotic Exit and Late-Born Cell Specification. Neuron 102, 1–16.

Cleary, M.D., and Doe, C.Q. (2006). Regulation of neuroblast competence: multiple temporal identity factors specify distinct neuronal fates within a single early competence window. Genes & development 20, 429–434.

Dall’Agnese, A., Caputo, L., Nicoletti, C., di Iulio, J., Schmitt, A., Gatto, S., Diao, Y., Ye, Z., Forcato, M., Perera, R., et al. (2019). Transcription Factor-Directed Re-wiring of Chromatin Architecture for Somatic Cell Nuclear Reprogramming toward trans-Differentiation. Mol Cell 76, 453–472 e458.

Davis, R.L., Weintraub, H., and Lassar, A.B. (1987). Expression of a single transfected cDNA converts fibroblasts to myoblasts. Cell 51, 987–1000.

De la Rossa, A., Bellone, C., Golding, B., Vitali, I., Moss, J., Toni, N., Luscher, C., and Jabaudon, D. (2013). In vivo reprogramming of circuit connectivity in postmitotic neocortical neurons. Nat Neurosci 16, 193–200.

de Melo, J., and Blackshaw, S. (2011). In vivo electroporation of developing mouse retina. J Vis Exp, 2847.

de Melo, J., Peng, G.H., Chen, S., and Blackshaw, S. (2011). The Spalt family transcription factor Sall3 regulates the development of cone photoreceptors and retinal horizontal interneurons. Development 138, 2325–2336.

Dong, X., Yang, H., Zhou, X., Xie, X., Yu, D., Guo, L., Xu, M., Zhang, W., Liang, G., and Gan, L. (2020). LIM-Homeodomain Transcription Factor LHX4 Is Required for the Differentiation of Retinal Rod Bipolar Cells and OFF-Cone Bipolar Subtypes. Cell reports 32, 108144.

Eden, E., Navon, R., Steinfeld, I., Lipson, D., and Yakhini, Z. (2009). GOrilla: a tool for discovery and visualization of enriched GO terms in ranked gene lists. BMC Bioinformatics 10, 48.

Elliott, J., Jolicoeur, C., Ramamurthy, V., and Cayouette, M. (2008). Ikaros confers early temporal competence to mouse retinal progenitor cells. Neuron 60, 26–39.

Elshatory, Y., Everhart, D., Deng, M., Xie, X., Barlow, R.B., and Gan, L. (2007). Islet-1 controls the differentiation of retinal bipolar and cholinergic amacrine cells. The Journal of neuroscience : the official journal of the Society for Neuroscience 27, 12707–12720.

Feng, J., Liu, T., Qin, B., Zhang, Y., and Liu, X.S. (2012). Identifying ChIP-seq enrichment using MACS. Nat Protoc 7, 1728–1740.

Furukawa, T., Morrow, E.M., and Cepko, C. (1997). Crx, a Novel otx-like Homeobox Gene, Shows Photoreceptor-Specific Expression and Regulates Photoreceptor Differentiation. Cell 91, 531–541.

Goldman, D. (2014). Muller glial cell reprogramming and retina regeneration. Nature reviews Neuroscience 15, 431–442.

Gotz, M., Sirko, S., Beckers, J., and Irmler, M. (2015). Reactive astrocytes as neural stem or progenitor cells: In vivo lineage, In vitro potential, and Genome-wide expression analysis. Glia 63, 1452–1468.

Grosskortenhaus, R., Pearson, B.J., Marusich, A., and Doe, C.Q. (2005). Regulation of temporal identity transitions in Drosophila neuroblasts. Developmental cell 8, 193–202.

Grosskortenhaus, R., Robinson, K.J., and Doe, C.Q. (2006). Pdm and Castor specify late-born motor neuron identity in the NB7-1 lineage. Genes & development 20, 2618–2627.

Gu, Z., Eils, R., and Schlesner, M. (2016). Complex heatmaps reveal patterns and correlations in multidimensional genomic data. Bioinformatics 32, 2847–2849.

Gurdon, J.B. (1962). The Developmental Capacity of Nuclei taken from Intestinal Epithelium Cells of Feeding Tadpoles. J Embryol exp Morph 10, 622–640.

Gurdon, J.B., and Melton, D.A. (2008). Nuclear Reprogramming in Cells. Science 322, 1811–1815.

Heizmann, B., Kastner, P., and Chan, S. (2017). The Ikaros family in lymphocyte development. Curr Opin Immunol 51, 14–23.

Hoang, T., Wang, J., Boyd, P., Wang, F., Santiago, C., Jiang, L., Yoo, S., Lahne, M., Todd, L.J., Jia, M., et al. (2020). Gene regulatory networks controlling vertebrate retinal regeneration. Science 370, eabb8598.

Holguera, I., and Desplan, C. (2018). Neuronal specification in space and time. Science 363, 176–180.

Honma, Y., Kiosawa, H., Mori, T., Oguri, A., Nikaido, T., Kanazawa, K., Tojo, M., Takeda, J., Tanno, Y., Yokoba, S., et al. (1999). Eos: a novel member of the Ikaros gene family expressed predominantly in the developing nervous system. FEBS letters 447, 76–80.

Isshiki, T., Pearson, B., Holbrook, S., and Doe, C.Q. (2001). Drosophila Neuroblasts Sequentially Express Transcription Factors which Specify the Temporal Identity of Their Neuronal Progeny. Cell 106, 511–521.

Iwafuchi-Doi, M., and Zaret, K.S. (2016). Cell fate control by pioneer transcription factors. Development 143, 1833–1837.

Jadhav, A.P., Roesch, K., and Cepko, C.L. (2009). Development and neurogenic potential of Muller glial cells in the vertebrate retina. Progress in retinal and eye research 28, 249–262.

Javed, A., Mattar, P., Lu, S., Kruczek, K., Kloc, M., Gonzalez-Cordero, A., Bremner, R., Ali, R.R., and Cayouette, M. (2020). Pou2f1 and Pou2f2 cooperate to control the timing of cone photoreceptor production in the developing mouse retina. Development 147, dev188730.

John, L.B., and Ward, A.C. (2011). The Ikaros gene family: transcriptional regulators of hematopoiesis and immunity. Mol Immunol 48, 1272–1278.

Jorstad, N.L., Wilken, M.S., Grimes, W.N., Wohl, S.G., VandenBosch, L.S., Yoshimatsu, T., Wong, R.O., Rieke, F., and Reh, T.A. (2017). Stimulation of functional neuronal regeneration from Muller glia in adult mice. Nature 548, 103–107.

Jozefczuk, J., Drews, K., and Adjaye, J. (2012). Preparation of mouse embryonic fibroblast cells suitable for culturing human embryonic and induced pluripotent stem cells. J Vis Exp, e3854.

Kambadur, R., Koizumi, K., Stivers, C., Nagle, J., Poole, S.J., and Odenwald, W.F. (1998). Regulation of POU genes by castor and hunchback establishes layered compartments in the Drosophila CNS. Genes & development 12, 246–260.

Kim, J.S., Sif, S., Jones, B., Jackson, A., Koipally, J., Heller, E., Winandy, S., Viel, A., Sawyer, A., Ikeda, T., et al. (1999). Ikaros DNA-Binding Proteins Direct Formation of Chromatin Remodeling Complexes in Lymphocytes. Immunity 10, 345–355.

Koche, R.P., Smith, Z.D., Adli, M., Gu, H., Ku, M., Gnirke, A., Bernstein, B.E., and Meissner, A. (2011). Reprogramming factor expression initiates widespread targeted chromatin remodeling. Cell Stem Cell 8, 96–105.

Kohwi, M., Lupton, J.R., Lai, S.L., Miller, M.R., and Doe, C.Q. (2013). Developmentally regulated subnuclear genome reorganization restricts neural progenitor competence in Drosophila. Cell 152, 97–108.

Lane, R.P., Cutforth, T., Young, J., Athanasiou, M., Friedman, C., Rowen, L., Evans, G., Axel, R., Hood, L., and Trask, B.J. (2001). Genomic analysis of orthologous mouse and human olfactory receptor loci. Proceedings of the National Academy of Sciences of the United States of America 98, 7390–7395.

Langmead, B., and Salzberg, S.L. (2012). Fast gapped-read alignment with Bowtie 2. Nature methods 9, 357–359.

Lee, Y., Messing, A., Su, M., and Brenner, M. (2008). GFAP promoter elements required for region-specific and astrocyte-specific expression. Glia 56, 481–493.

Lehre, K.P., Davanger, S., and Danbolt, N.C. (1997). Localization of the glutamate transporter protein GLAST in rat retina. Brain research 744, 129–137.

Lenkowski, J.R., and Raymond, P.A. (2014). Muller glia: Stem cells for generation and regeneration of retinal neurons in teleost fish. Progress in retinal and eye research 40, 94–123.

Li, H., and Chen, G. (2016). In Vivo Reprogramming for CNS Repair: Regenerating Neurons from Endogenous Glial Cells. Neuron 91, 728–738.

Liu, Y., Yu, C., Daley, T.P., Wang, F., Cao, W.S., Bhate, S., Lin, X., Still, C., 2nd, Liu, H., Zhao, D., et al. (2018). CRISPR Activation Screens Systematically Identify Factors that Drive Neuronal Fate and Reprogramming. Cell Stem Cell 23, 758–771 e758.

Love, M.I., Huber, W., and Anders, S. (2014). Moderated estimation of fold change and dispersion for RNA-seq data with DESeq2. Genome Biol 15, 550.

Mall, M., Kareta, M.S., Chanda, S., Ahlenius, H., Perotti, N., Zhou, B., Grieder, S.D., Ge, X., Drake, S., Euong Ang, C., et al. (2017). Myt1l safeguards neuronal identity by actively repressing many non-neuronal fates. Nature 544, 245–249.

Martin, J.F., and Poche, R.A. (2019). Awakening the regenerative potential of the mammalian retina. Development 146.

Matsuda, T., and Cepko, C.L. (2004). Electroporation and RNA interference in the rodent retina in vivo and in vitro. Proceedings of the National Academy of Sciences of the United States of America 101, 16–22.

Mattar, P., Ericson, J., Blackshaw, S., and Cayouette, M. (2015). A conserved regulatory logic controls temporal identity in mouse neural progenitors. Neuron 85, 497–504.

Mattar, P., Jolicoeur, C., Dang, T., Shah, S., Clark, B.S., and Cayouette, M. (2021). A Casz1-NuRD complex regulates temporal identity transitions in neural progenitors. Scientific reports 11, 3858.

Mayran, A., Khetchoumian, K., Hariri, F., Pastinen, T., Gauthier, Y., Balsalobre, A., and Drouin, J. (2018). Pioneer factor Pax7 deploys a stable enhancer repertoire for specification of cell fate. Nat Genet 50, 259–269.

McLean, C.Y., Bristor, D., Hiller, M., Clarke, S.L., Schaar, B.T., Lowe, C.B., Wenger, A.M., and Bejerano, G. (2010). GREAT improves functional interpretation of cis-regulatory regions. Nat Biotechnol 28, 495–501.

Mor, N., Rais, Y., Sheban, D., Peles, S., Aguilera-Castrejon, A., Zviran, A., Elinger, D., Viukov, S., Geula, S., Krupalnik, V., et al. (2018). Neutralizing Gatad2a-Chd4- Mbd3/NuRD Complex Facilitates Deterministic Induction of Naive Pluripotency. Cell Stem Cell 23, 412–425 e410.

Nathans, J. (2010). Generation of an inducible Slc1a3-cre/ERT transgenic allele. MGI Direct Data Submission MGI: J:157151.

Novotny, T., Eiselt, R., and Urban, J. (2002). Hunchback is required for the specification of the early sublineage of neuroblast 7-3 in the *Drosophila* central nervous system. Development 129, 1027–1036.

O’Neill, D.W., Schoetz, S.S., Lopez, R.A., Castle, M., Rabinowitz, L., Shor, E., Krawchik, D., Goll, M.G., Renz, M., Seelig, H.-P., et al. (2000). An Ikaros-Containing Chromatin-Remodeling Complex in Adult-Type Erythroid Cells. Molecular and cellular biology 20, 7572–7582.

Oberst, P., Agirman, G., and Jabaudon, D. (2019). Principles of progenitor temporal patterning in the developing invertebrate and vertebrate nervous system. Curr Opin Neurobiol 56, 185–193.

Ortin-Martinez, A., Tsai, E.L., Nickerson, P.E., Bergeret, M., Lu, Y., Smiley, S., Comanita, L., and Wallace, V.A. (2016). A Reinterpretation of Cell Transplantation: GFP Transfer from Donor to Host Photoreceptors. Stem cells 35, 932–939.

Patro, R., Duggal, G., Love, M.I., Irizarry, R.A., and Kingsford, C. (2017). Salmon provides fast and bias-aware quantification of transcript expression. Nature methods 14, 417–419.

Pearson, B.J., and Doe, C.Q. (2003). Regulation of neuroblast competence in Drosophila. Nature 425, 624–628.

Pearson, R.A., Gonzalez-Cordero, A., West, E.L., Ribeiro, J.R., Aghaizu, N., Goh, D., Sampson, R.D., Georgiadis, A., Waldron, P.V., Duran, Y., et al. (2016). Donor and host photoreceptors engage in material transfer following transplantation of post-mitotic photoreceptor precursors. Nature communications 7, 13029.

Plessy, C., Pascarella, G., Bertin, N., Akalin, A., Carrieri, C., Vassalli, A., Lazarevic, D., Severin, J., Vlachouli, C., Simone, R., et al. (2012). Promoter architecture of mouse olfactory receptor genes. Genome Res 22, 486–497.

Pollak, J., Wilken, M.S., Ueki, Y., Cox, K.E., Sullivan, J.M., Taylor, R.J., Levine, E.M., and Reh, T.A. (2013). ASCL1 reprograms mouse Muller glia into neurogenic retinal progenitors. Development 140, 2619–2631.

Powell, C., Elsaeidi, F., and Goldman, D. (2012). Injury-dependent Muller glia and ganglion cell reprogramming during tissue regeneration requires Apobec2a and Apobec2b. The Journal of neuroscience : the official journal of the Society for Neuroscience 32, 1096–1109.

Rais, Y., Zviran, A., Geula, S., Gafni, O., Chomsky, E., Viukov, S., Mansour, A.A., Caspi, I., Krupalnik, V., Zerbib, M., et al. (2013). Deterministic direct reprogramming of somatic cells to pluripotency. Nature 502, 65–70.

Ramirez, F., Ryan, D.P., Gruning, B., Bhardwaj, V., Kilpert, F., Richter, A.S., Heyne, S., Dundar, F., and Manke, T. (2016). deepTools2: a next generation web server for deep-sequencing data analysis. Nucleic acids research 44, W160–165.

Reichenbach, A., and Bringmann, A. (2013). New functions of Muller cells. Glia 61, 651–678.

Robinson, J.T., Thorvaldsdóttir, H., Winckler, W., Guttman, M., Lander, E.S., Getz, G., and Mesirov, J.P. (2011). Integrative genomics viewer. Nat Biotechnol 29, 24–26.

Roesch, K., Jadhav, A.P., Trimarchi, J.M., Stadler, M.B., Roska, B., Sun, B.B., and Cepko, C.L. (2008). The transcriptome of retinal Muller glial cells. The Journal of comparative neurology 509, 225–238.

Rossi, A.M., Fernandes, V.M., and Desplan, C. (2017). Timing temporal transitions during brain development. Curr Opin Neurobiol 42, 84–92.

Rouaux, C., and Arlotta, P. (2013). Direct lineage reprogramming of post-mitotic callosal neurons into corticofugal neurons in vivo. Nature cell biology 15, 214–221.

Santos-Ferreira, T., Llonch, S., Borsch, O., Postel, K., Haas, J., and Ader, M. (2016). Retinal transplantation of photoreceptors results in donor–host cytoplasmic exchange. Nature communications 7, 13028.

Sapkota, D., Chintala, H., Wu, F., Fliesler, S.J., Hu, Z., and Mu, X. (2014). Onecut1 and Onecut2 redundantly regulate early retinal cell fates during development. Proceedings of the National Academy of Sciences of the United States of America 111, E4086–4095.

Sen, S.Q., Chanchani, S., Southall, T.D., and Doe, C.Q. (2019). Neuroblast-specific open chromatin allows the temporal transcription factor, Hunchback, to bind neuroblast-specific loci. eLife 8, e44036.

Shekhar, K., Lapan, S.W., Whitney, I.E., Tran, N.M., Macosko, E.Z., Kowalczyk, M., Adiconis, X., Levin, J.Z., Nemesh, J., Goldman, M., et al. (2016). Comprehensive Classification of Retinal Bipolar Neurons by Single-Cell Transcriptomics. Cell 166, 1308–1323 e1330.

Singh, M.S., Balmer, J., Barnard, A.R., Aslam, S.A., Moralli, D., Green, C.M., Barnea-Cramer, A., Duncan, I., and MacLaren, R.E. (2016). Transplanted photoreceptor precursors transfer proteins to host photoreceptors by a mechanism of cytoplasmic fusion. Nature communications 7, 13537.

Su, M., Hu, H., Lee, Y., d’Azzo, A., Messing, A., and Brenner, M. (2004). Expression Specificity of GFAP Transgenes. Neurochemical Research 29, 2075–2093.

Supek, F., Bosnjak, M., Skunca, N., and Smuc, T. (2011). REVIGO summarizes and visualizes long lists of gene ontology terms. PloS one 6, e21800.

Takahashi, K., and Yamanaka, S. (2006). Induction of pluripotent stem cells from mouse embryonic and adult fibroblast cultures by defined factors. Cell 126, 663–676.

Tsunemoto, R., Lee, S., Szucs, A., Chubukov, P., Sokolova, I., Blanchard, J.W., Eade, K.T., Bruggemann, J., Wu, C., Torkamani, A., et al. (2018). Diverse reprogramming codes for neuronal identity. Nature 557, 375–380.

Ueki, Y., Wilken, M.S., Cox, K.E., Chipman, L., Jorstad, N., Sternhagen, K., Simic, M., Ullom, K., Nakafuku, M., and Reh, T.A. (2015). Transgenic expression of the proneural transcription factor Ascl1 in Muller glia stimulates retinal regeneration in young mice. Proceedings of the National Academy of Sciences of the United States of America 112, 13717–13722.

Van de Sande, B., Flerin, C., Davie, K., De Waegeneer, M., Hulselmans, G., Aibar, S., Seurinck, R., Saelens, W., Cannoodt, R., Rouchon, Q., et al. (2020). A scalable SCENIC workflow for single-cell gene regulatory network analysis. Nat Protoc 15, 2247–2276.

van der Raadt, J., van Gestel, S.H.C., Nadif Kasri, N., and Albers, C.A. (2019). ONECUT transcription factors induce neuronal characteristics and remodel chromatin accessibility. Nucleic acids research 47, 5587–5602.

Vierbuchen, T., Ostermeier, A., Pang, Z.P., Kokubu, Y., Sudhof, T.C., and Wernig, M. (2010). Direct conversion of fibroblasts to functional neurons by defined factors. Nature 463, 1035–1041.

Wang, L.-L., Garcia, C.S., Zhong, X., Ma, S., and Zhang, C.-L. (2020). Rapid and efficient in vivo astrocyte-to-neuron conversion with regional identity and connectivity? bioRxiv.

Wapinski, O.L., Lee, Q.Y., Chen, A.C., Li, R., Corces, M.R., Ang, C.E., Treutlein, B., Xiang, C., Baubet, V., Suchy, F.P., et al. (2017). Rapid Chromatin Switch in the Direct Reprogramming of Fibroblasts to Neurons. Cell reports 20, 3236–3247.

Wapinski, O.L., Vierbuchen, T., Qu, K., Lee, Q.Y., Chanda, S., Fuentes, D.R., Giresi, P.G., Ng, Y.H., Marro, S., Neff, N.F., et al. (2013). Hierarchical mechanisms for direct reprogramming of fibroblasts to neurons. Cell 155, 621–635.

Yao, K., Qiu, S., Wang, Y.V., Park, S.J.H., Mohns, E.J., Mehta, B., Liu, X., Chang, B., Zenisek, D., Crair, M.C., et al. (2018). Restoration of vision after de novo genesis of rod photoreceptors in mammalian retinas. Nature 560, 484–488.

Zhou, H., Su, J., Hu, X., Zhou, C., Li, H., Chen, Z., Xiao, Q., Wang, B., Wu, W., Sun, Y., et al. (2020). Glia-to-Neuron Conversion by CRISPR-CasRx Alleviates Symptoms of Neurological Disease in Mice. Cell.

