## Supplemental Figures and Tables for "Direct neuronal reprogramming by temporal identity factors"

**SUPPLEMENTARY FIGURES AND TABLES**

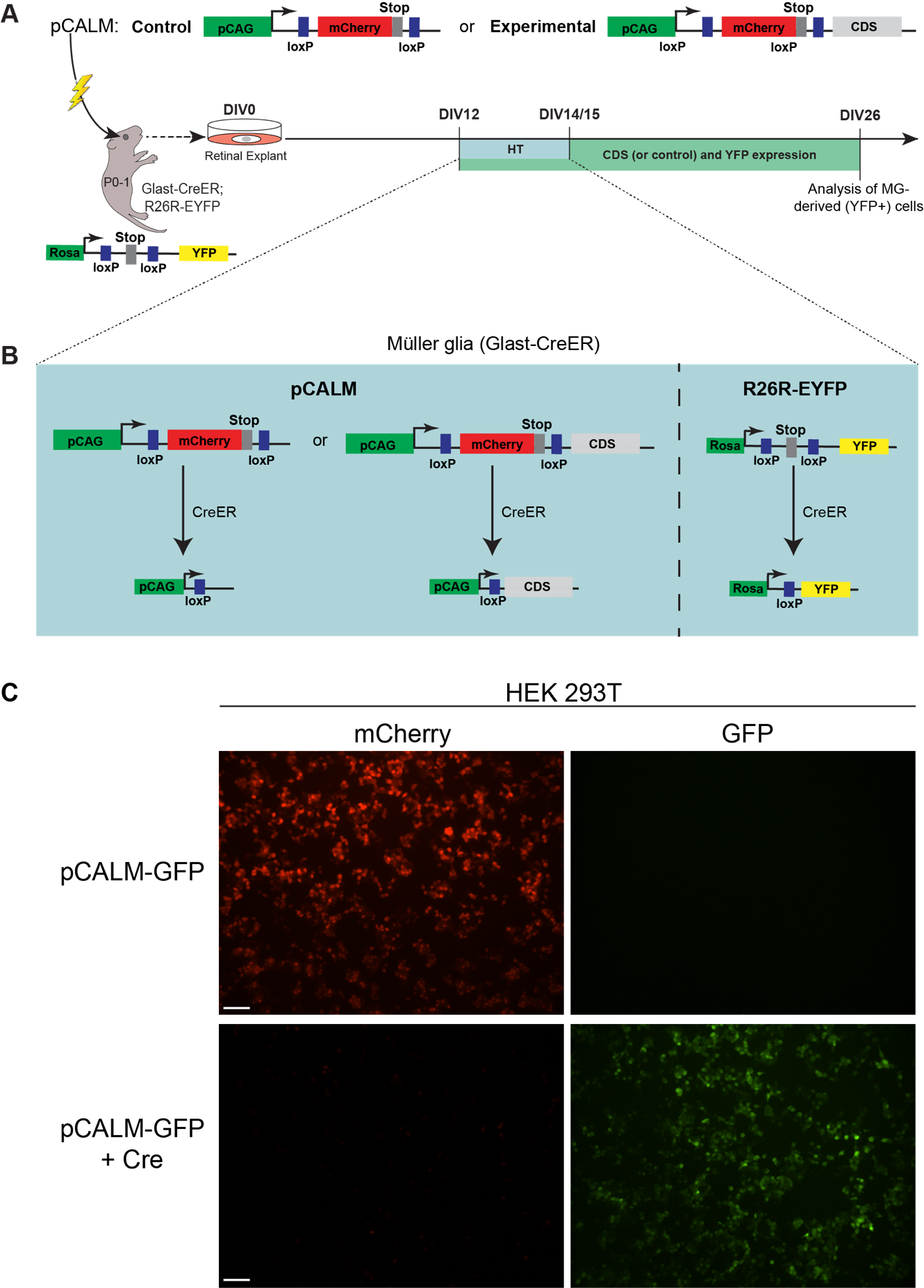

**Figure S1: Detailed representation of experimental procedure. Related to Figure 1.**

**(A)** pCALM plasmids, control or experimental, are electroporated (lightning bolt) in the eyes of neonate Glast-CreER;RosaYFP animals, which are explanted and cultured for 26 days. Hydroxytamoxifen (HT) is added to culture medium from DIV12 to 14/15. CDS: Coding sequence. **(B)** Representation of CreER activity in Müller glia when HT is present. pCALM constructs (left) are recombined at the loxP cassettes to induce expression of the gene coding sequence (experimental) or nothing (control). Rosa locus (right) loxP cassette is also removed to induce permanent expression of EYFP. **(C)** HEK 293T cells transfected with pCALM-GFP (top row) or pCALM-GFP + Cre (bottom row). Scale bars: 100µm.

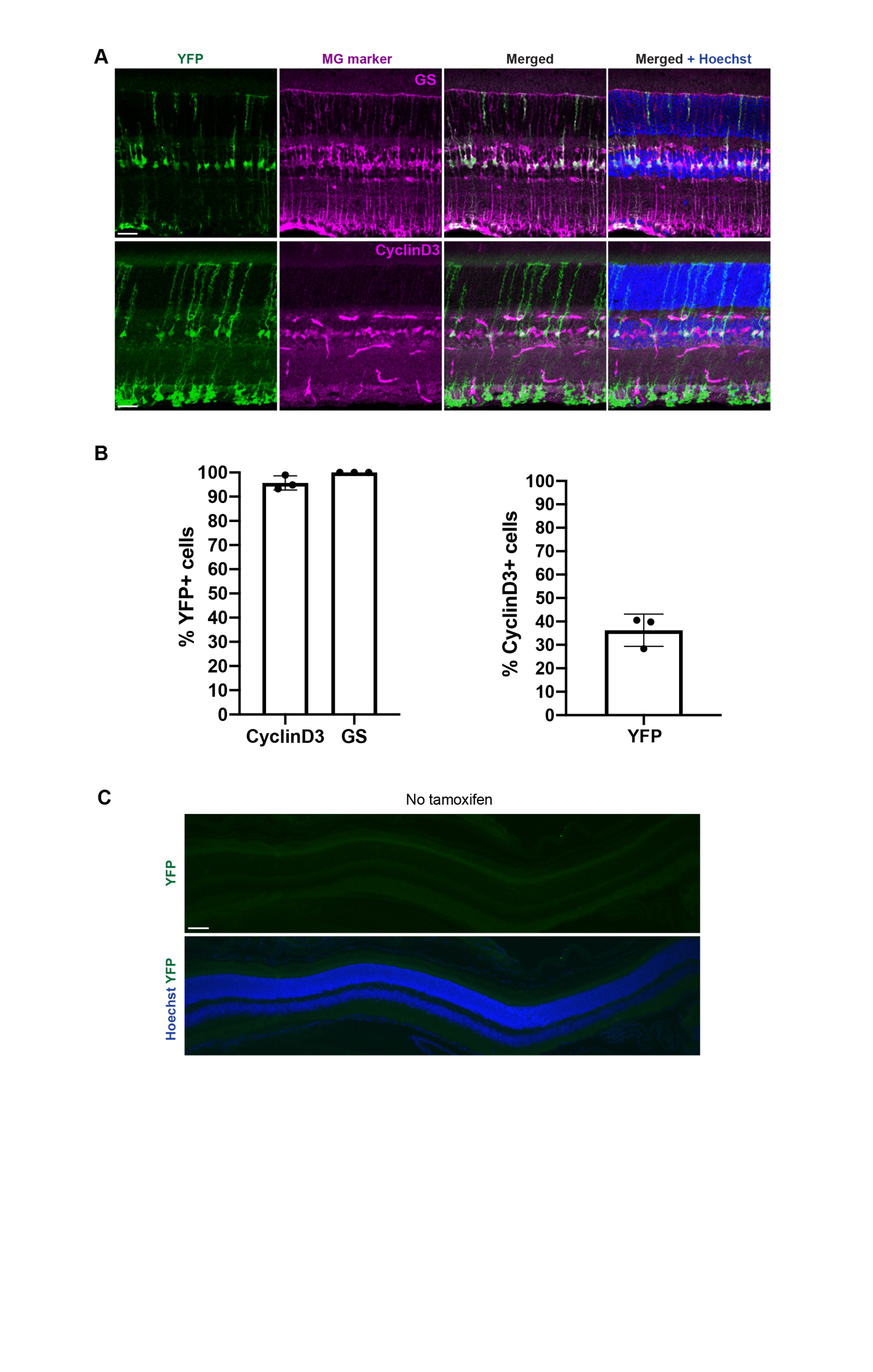

**Figure S2: Glast-CreER;R26R-EYFP mouse line specifically labels MG. Related to Figure 1.**

**(A)** Co-immunostaining for YFP and glutamine synthetase (GS; top) or CyclinD3 (bottom) on retinal sections from the Glast-CreER;R26R-EYFP mouse line 7-8 days after tamoxifen injection. Scale bars: 25µm. **(B)** Left: Graph representing the proportion of YFP+ cells co-labelled with MG markers. Right: Proportion of CyclinD3+ cells co-labelled with YFP. Quantifications done in the Glast-CreER;R26R-EYFP mouse line 7-8 days after tamoxifen injection. **(C)** Representative images of retinal sections, immunostained for YFP, of CreER;R26R-EYFP mice that did not received tamoxifen. Scale bars: 50µm. (A;top) shows single plane images, whereas (A;bottom) and (E) are z projections. Graphs represent mean +/- standard deviation.

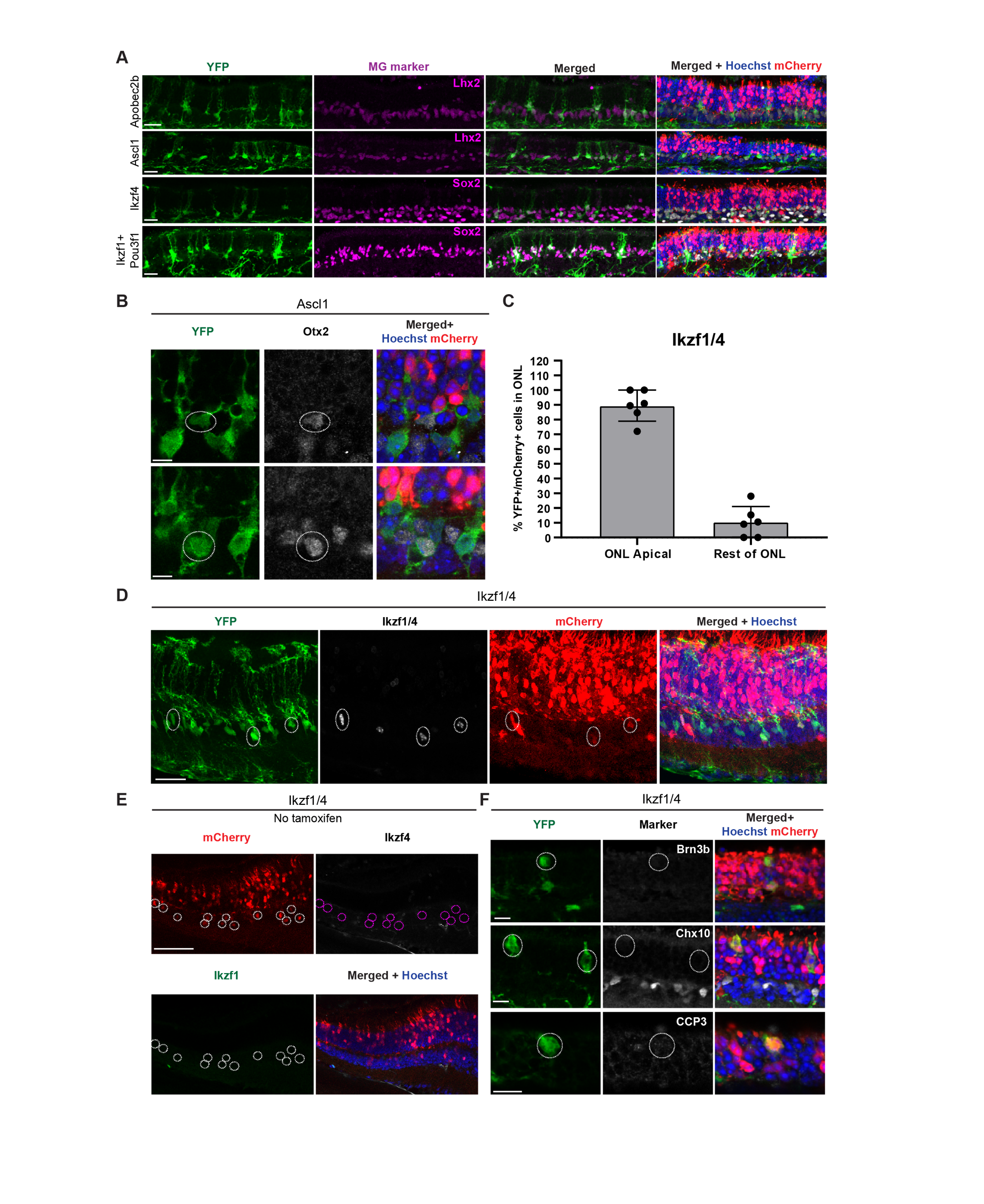

**Figure S3: Complementary data from the MG reprogramming ex vivo screen. Related to Figures 1 and 2.**

**(A)** Representative images of co-immunostaining for YFP and the MG markers Lhx2 (top) and Sox2 (bottom) on retinal sections from additional conditions tested in the screen. Scale bars: 24µm. **(B)** Co-immunostaining for YFP and Otx2 after electroporation of Ascl1. Circles show co-labelled cells. Scale bars: 7µm. **(C)** Quantification of the localization of YFP+/mCherry+ morphologically reprogrammed cells in the ONL in the Ikzf1/4 condition. Graph shows mean +/- standard deviation. **(D)** Co-immunostaining for YFP and Ikzf1/4 on section of Ikzf1/4 in vivo electroporated retinas 5 weeks post-tamoxifen. Circles show YFP+/mCherry+/Ikzf1/4+ cells. Scale bar: 25 µm. **(E)** Co-immunostaining for Ikzf4 and Ikzf1 on a retinal section of Ikzf1/4 in vivo electroporated retinas 6.5 weeks post-electroporation without tamoxifen. Circles point to INL mCherry+ cells. Scale bar: 60µm. **(F)** Co-immunostaining for YFP and Brn3b, Chx10, and cleaved caspase-3 (CCP3) in Ikzf1/4-electroporated retinas. Reprogrammed cells (circled) were negative for all three markers. Scale bars: 12µm. (A) shows z-projections, all other images are single planes.

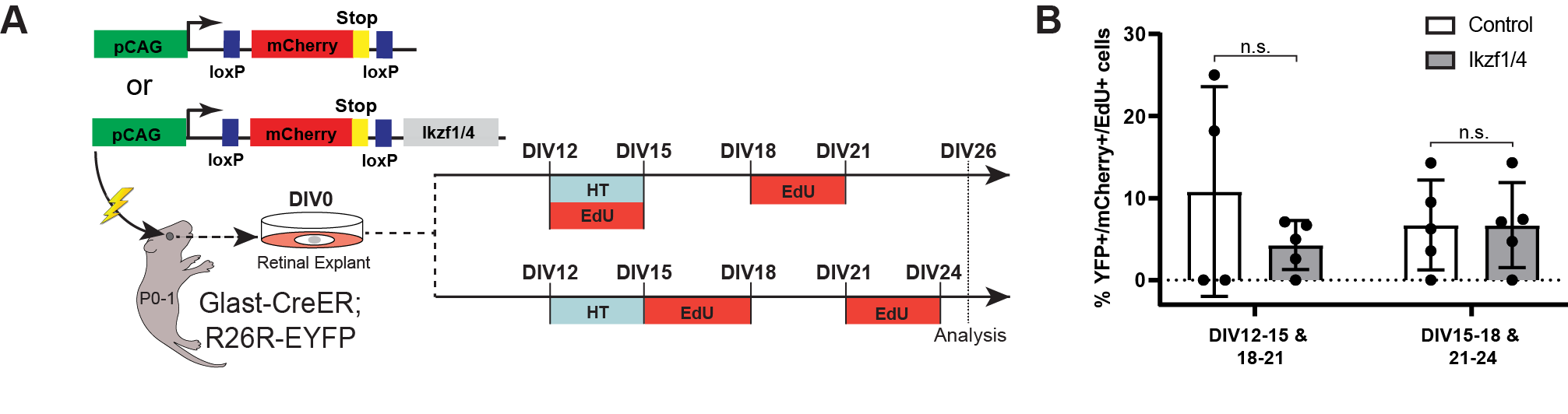

**Figure S4: Ikzf1/4 do not promote MG cell cycle re-entry. Related to Figures 1 and 2.**

**(A)** Schematic representation of the cell cycle re-entry analysis protocol. DIV: days in vitro, HT: hydroxy-tamoxifen. CDS: coding sequence. Lightning bolt represents electroporation. **(B)** Quantifications of the number of YFP+/mCherry+ cells that incorporated EdU in control and Ikzf1/4-electroporated retinas in both conditions. Graph shows mean +/- standard deviation. n.s.: not significant; unpaired t-tests, n=4 in control DIV12-15, n=5 in control DIV15-18, n=5 in both Ikzf1/4 conditions.

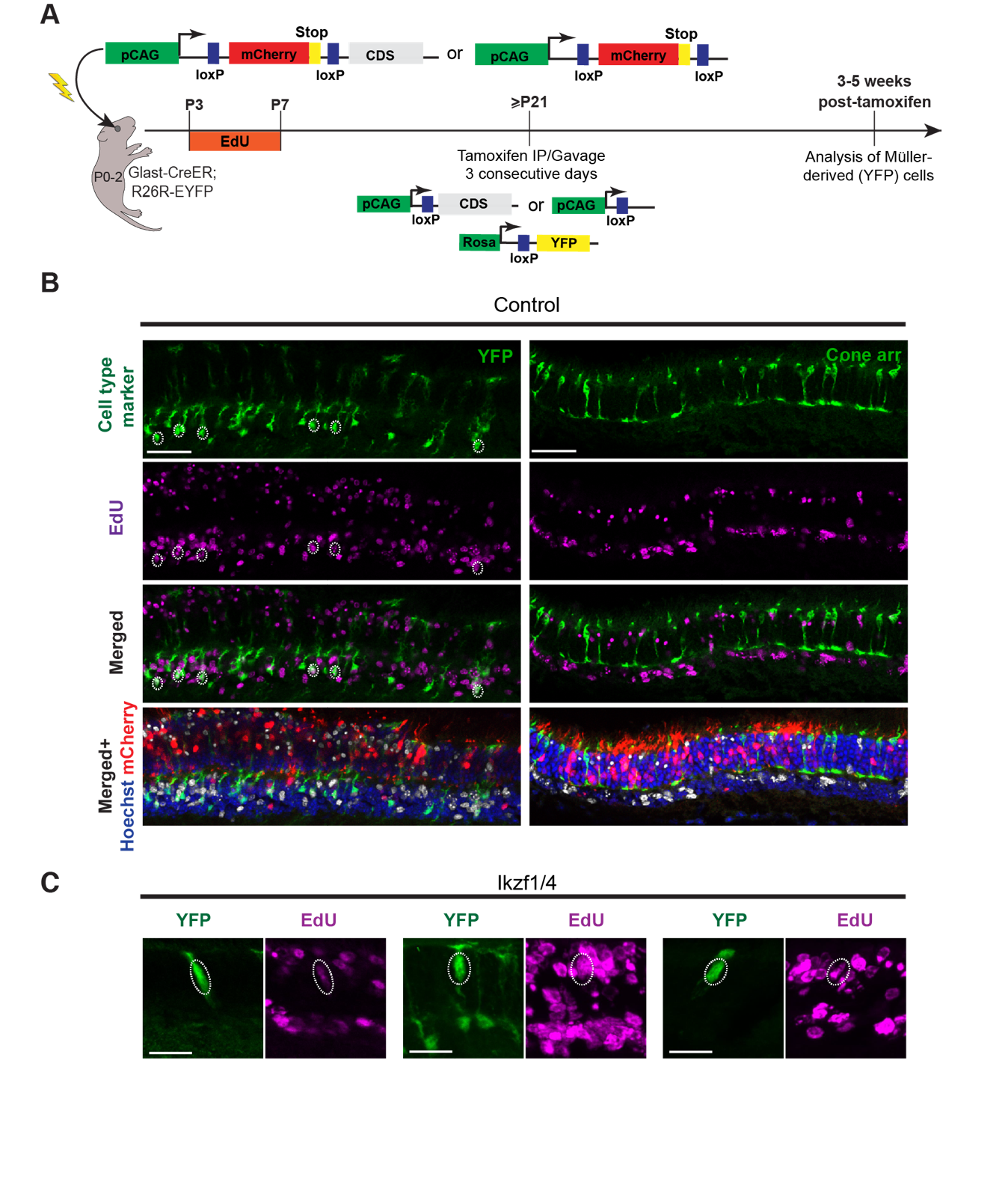

**Figure S5: Detection of reprogrammed cells is not the result of material exchange. Related to Figure 3.**

**(A)** Schematic representation of the experimental protocol. IP: intraperitoneal injection. Lightning bolt represents electroporation. **(B)** Co-staining for EdU and YFP (left) or cone arrestin (cone arr.; right) on retinal sections from control-electroporated retinas. YFP+ MG (circled) stained for EdU, whereas cone arrestin+ cone photoreceptors did not (n=3). Scale bars: 33µm. **(C)** Co-staining for EdU and YFP on retinal sections after Ikzf1/4 electroporation. Reprogrammed cells (circled) in the ONL co-label with EdU. Scale bars: 15µm. All images are single planes except (C; middle and right), which are z projections.

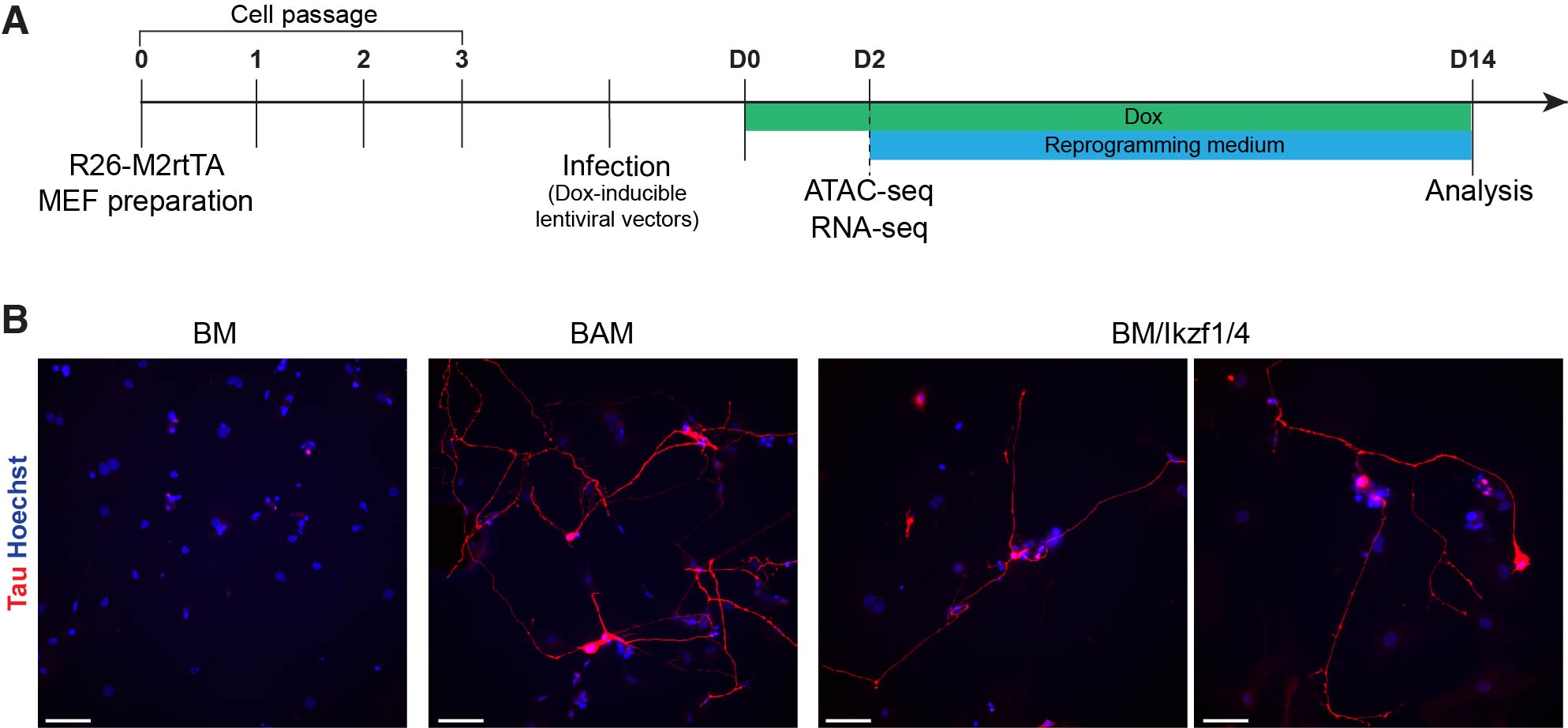

**Figure S6: BM/Ikzf1/4 convert MEFs into iNs. Related to Figure 6.**

**(A)** Schematic representation of the MEF reprogramming protocol. Day 0 (D0) of reprogramming assay corresponds to the first day of doxycycline-induced (dox) expression of lentiviral vectors. **(B)** Representative images of MEF cultures expressing BM, BAM, or BM/Ikzf1/4 at D14 of reprogramming assay immunostained for the neuronal marker Tau. Scale bars: 50µm.

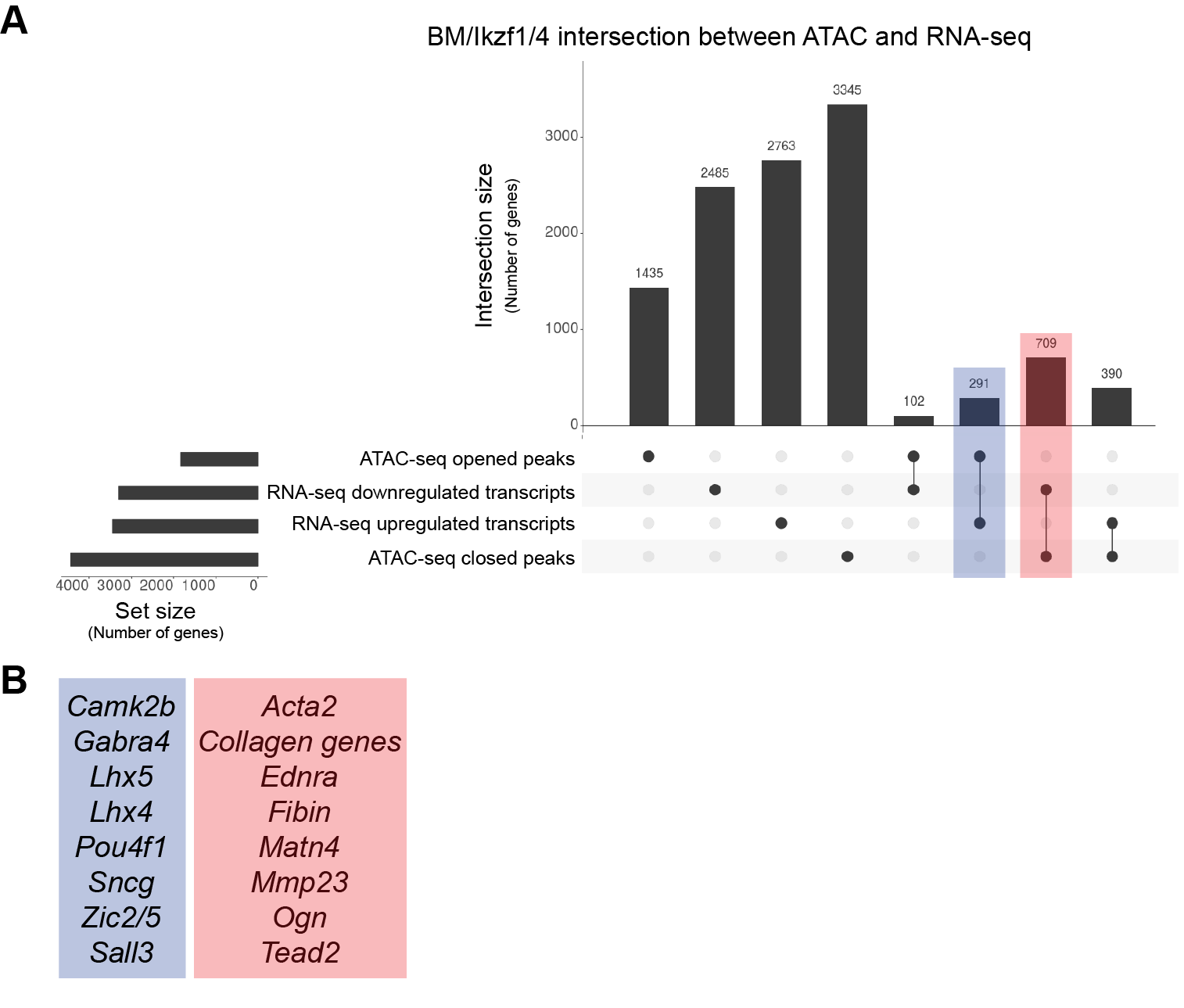

**Figure S7: Correlation between ATAC and RNA-sequencing data. Related to Figure 6.**

**(A)** Upset plot showing correlation between ATAC and RNA-seq hits for BM/Ikzf1/4 in MEFs. Y-axis represents intersection size as number of genes, and x-axis represents conditions (black dots indicating condition with connected black dots indicating intersection). Set size represents the total number of genes associated with each condition. ATAC-seq opened and closed genes correspond to genes associated with peaks +/- 2kb from TSS. Blue highlights intersection of genes associated with opened BM/Ikzf1/4 peaks and BM/Ikzf1/4 upregulated transcripts. Red highlights intersection of genes associated with closed BM/Ikzf1/4 peaks and BM/Ikzf1/4 downregulated transcripts. **(B)** Example of genes included in both intersections. Colors correspond to (A). See Table S4 for full list.

| Cell types labelled | Markers screened | Presence in reprogrammed cells |
| --- | --- | --- |
| Müller glia | Lhx2 | Few cells |
| Müller glia | Sox2 | Few cells |
| Müller glia | CyclinD3 | Few cells |
| Cone photoreceptors and retinal ganglion cells | Rxrg | Yes |
| Cone photoreceptors | S-opsin | No |
| Cone photoreceptors | L/M-opsin | No |
| Cone photoreceptors | Cone arrestin | No |
| Cone photoreceptors | PNA | No |
| Rod photoreceptors | Nrl | No |
| Photoreceptors and bipolar cells | Otx2 | No |
| Bipolar cells | Chx10 | No |
| Retinal ganglion cells | Brn3b | No |
| Apoptosis | Cleaved caspase 3 | No |

**Table S1: Summary of cell-type specific marker expression screened by immunostaining in Ikzf1/4-electroporated condition ex vivo. Related to Figure 2.**

| Antigen | Species | Company | Cat. number | Concentration |
| --- | --- | --- | --- | --- |
| Brn3b | Goat | Santa Cruz Biotechnology | SC-6026 | 1/200 |
| Chx10 | Sheep | Exalpha Biologicals | X1180P | 1/500 |
| Cleaved caspase 3 | Rabbit | New England Biolabs | 9661 | 1/100 |
| Cone arrestin | Rabbit | Millipore Sigma | AB15282 | 1/1000 |
| CyclinD3 | Mouse | Santa Cruz Biotechnology | SC-6283 | 1/100 |
| GFP | Chicken | Abcam | AB13970 | 1/1000 |
| GFP | Rabbit | Thermo Fisher | A11122 | 1/500 |
| Glutamine Synthetase | Mouse | BD Bioscience | 610517 | 1/200 |
| Glutamine Synthetase | Mouse | Chemicon | MAB-302a | 1/200 |
| Ikzf1 | Goat | Santa Cruz Biotechnology | discontinued | 1/200 |
| Ikzf4 | Mouse | Sigma | SAB1407877 | 1/500 |
| Ki-67 | Rabbit | Neo Markers | RM-9106-S1 | 1/200 |
| L/M-opsin | Rabbit | Millipore Sigma | AB5405 | 1/1000 |
| Lhx2 | Rabbit | Thermo Fisher | PA5-78287 | 1/200 |
| Nrl | Goat | R&D Systems | AF2945-SP | 1/500 |
| Otx2 | Goat | R&D Systems | AF1979 | 1/1000 |
| Pax6 | Rabbit | Millipore Sigma | AB2237 | 1/200 |
| PNA (Lectin conjugates) | N/A | Molecular Probes | L-32460 | 1/500 |
| Rxrg | Rabbit | Abcam | AB15518 | 1/200 |
| S-opsin | Goat | Santa Cruz Biotechnology | SC-14363P | 1/1000 |
| Sox2 | Rabbit | Abcam | AB97959 | 1/200 |
| Tau | Rabbit | Dako | M4403 | 1/10000 |

**Methods Table S1: Primary antibodies. Related to *Immunostaining* section of methods.**

**OTHER SUPPLEMENTARY MATERIAL**

**Table S2: Significantly enriched and depleted ATAC peaks +/- 2kb from TSS in MEFs 48 hours after expression of BM/Ikzf1/4 compared to BM. Related to Figure 6.**

**Table S3: RNA-sequencing data of MEFs 48 hours after expression of BM/Ikzf1/4 compared to BM. Related to Figure 6.**

**Table S4: BM/Ikzf1/4 ATAC and RNA-sequencing intersection data of MEFs 48 hours after of expression BM/Ikzf1/4. Related to Figures 6 and S7.**

**Video S1: MEF expressing BM/Ikzf1 reprogram to iN without proliferating. Related to Figures 6 and S6.**

Time-lapse recording of MEFs isolated from Tau::EGFP mice transfected with BM/Ikzf1. GFP signal reveals Tau expression. Note the upregulation of Tau::EGFP and drastic morphological change occurring over time. Photos taken every 8 hours.
